# Mass Generation and Long-term Expansion of Hepatobiliary Organoids from Adult Primary Human Hepatocytes

**DOI:** 10.1101/2024.06.10.598262

**Authors:** Ary Marsee, Arabela Ritchie, Adam Myszczyszyn, Shicheng Ye, Jung-Chin Chang, Arif Ibrahim Ardisasmita, Indi P Joore, Jose Castro-Alpízar, Sabine A Fuchs, Kerstin Schneeberger, Bart Spee

## Abstract

Adult primary human hepatocytes (PHHs) are the gold standard in *ex vivo* toxicological studies and possess the clinical potential to treat patients with liver disease as advanced therapy medicinal products (ATMPs). However, the utility of this valuable cell type has been limited by short-term functionality and limited expansion potential *in vitro*. While notable advances have been made in the long-term maintenance of primary hepatocytes, there has been limited success in driving the efficient generation and expansion of adult PHH-derived organoids which recapitulate both liver tissue architecture and function, hampering *in vitro* studies and regenerative medicine applications. Here we describe the mass generation and long-term expansion of hepatobiliary organoids with functionally interconnected hepatic and biliary-like structures from adult primary human hepatocytes. Hepatobiliary organoids retain the expression of lineage and functional markers, closely resembling PHH, while also acquiring the expression of regeneration, fetal and biliary markers. Organoids perform key hepatocyte functions while proliferating and can be matured to enhance their functionality. As a proof-of-principle, we demonstrate that hepatobiliary organoids can recapitulate hallmarks of cholestasis and steatosis *in vitro*. Moreover, we show that hepatocytes can be transfected, transduced and gene edited in 3D prior to organoid generation, facilitating a wide range of applications. Our novel hepatobiliary organoid system bridges the gap between short-term functionality of primary human hepatocytes and the need for scalable, long-term organoid models of the adult liver, offering immense potential for drug testing, disease modeling, and advanced therapeutic applications.

## Introduction

The liver is an essential organ responsible for a multitude of vital functions, from the maintenance of lipid and glucose homeostasis, to the production of serum proteins and the detoxification of xenobiotics. Hepatocytes, the parenchymal cells of the liver which make up 80% of its mass, are responsible for most of these functions^1^. The ability to maintain adult primary human hepatocytes (PHHs) *in vitro* has enabled applications ranging from pharmacological and toxicological testing^2^, to regenerative medicine as advanced therapy medicinal products (ATMPs)^3^. However, primary hepatocytes have proven to be a fastidious cell type, limiting their translational utility *in vitro* and clinical application *in vivo*. Under traditional hepatocytes culture are conditions, primary phenotypically unstable, dedifferentiating within hours to days^4^. As this occurs, cultured hepatocytes progressively lose their morphology, polarity, and functionality^5^. Several techniques have been developed to prolong the functionality of primary hepatocytes *in vitro*, such as sandwich^6^ and spheroid^7^ culture, which preserve the hepatocyte phenotype for 1-2 months^8,9^. However, while these techniques have extended the culturable period of hepatocytes, there has been limited success in generating three dimensional (3D) models which recapitulate both liver tissue architecture and function.

While spheroid culture of PHHs enables long-term studies on functional hepatocytes, such as repeated dosing of potentially toxic compounds, their simplistic organization is distinct to that of the hepatocytes *in situ*. For example, in the liver hepatocytes are arranged in cords, spanning the portal-central axis, with their basolateral membrane facing the space of Disse and incoming blood supply, and their apical membrane forming bile canaliculi between adjacent hepatocytes^10^. Typically, hepatocyte cords in the homeostatic adult liver are 1-2 hepatocytes in thickness, maximizing the surface area available to perform the diverse array of metabolic functions hepatocytes are responsible for^11^. This is in stark contrast to the dense, cell packed organization of spheroids, which only directly expose hepatocytes on the periphery to the outside environment. Furthermore, the (partial) localization of canalicular transporters to the exterior^12^ indicates that hepatocyte spheroids expose portions of their apical membrane outward, facing the medium (Figure S1). In normal liver architecture the apical membrane of hepatocytes is sequestered to regions between hepatocytes, and thus not facing the incoming blood supply^13^. Finally, in the liver hepatocyte cords terminate in bile ducts, which transport bile secreted by hepatocytes. Current spheroid and organoid models derived from adult PHHs lack this hepato-biliary connection.

In addition to challenges in recapitulating the complex structure of the native liver, driving the expansion of adult PHH while preserving their functionality and identity has also proven difficult. Recent studies have reported the expansion of human hepatocytes in 2D as de-differentiated progenitor-like cells, expressing biliary/stem cell markers *EPCAM, KRT7/19* and *SOX9*, and their subsequent re-differentiation towards the hepatocyte lineage. However, these cells do not fully reacquire their hepatocyte morphology and mature phenotype after re-differentiation^14–17^. Breakthroughs in the long-term expansion of functional primary hepatocytes, without reversion to a progenitor cell type, were reported in 2018^18,19^. Utilizing TNFα in combination with a cocktail of growth factors and small molecules, Peng *et al*. described the clonal expansion of single mouse hepatocytes for more than 8 months^18^. Using a similar approach, Hu *et al*. described the expansion of mouse hepatocytes (2-3 months), as well as human fetal liver cells (>11 months), as epithelial organoids. The group also reported the establishment of pediatric and adult human hepatocyte organoids (HOs). However, organoid formation efficiency was low, with 0.5-1% of plated adult human hepatocytes generating organoids with limited expansion potential^19^. Recently, Hendricks *et al*. demonstrated that HOs generated under the conditions described by Hu *et al*. struggled to exceed a diameter of 40 µm over a 14-day period. To facilitate expansion of cultures the media was supplemented with IL-6 and an FXR agonist, which promoted the expansion of PHH, with neonatal and infant hepatocytes expanded up to 4 months^20^. In 2020, Lauschke and colleagues demonstrated that activation of Wnt/β-catenin signaling during aggregation of PHHs stimulated a proliferative response, consistent with the literature on hepatocyte regeneration *in vivo*^21^. Building on this work, the group reported the induction of PHH proliferation in hepatocyte spheroids through the synergistic activation of Wnt/β-catenin and NFκB signaling in growth factor-free conditions^22^. In a separate study, Rose *et al*. reported that aggregation of PHH followed by transfer to a collagen gel stimulated their expansion, with the group describing two waves of hepatocyte proliferation, wherein hepatocytes were organized as hollow spheroids^23^.

Here we describe a novel method that enables the mass generation and long-term expansion (2-3 months) of functional hepatobiliary organoids (HBOs) derived from cryopreserved adult primary human hepatocytes. Rather than capturing the low percentage of hepatocytes amenable to single-cell expansion, our method captures the broader hepatocyte population, enabling the mass generation of expanding organoids with interconnected hepatic and biliary-like structures from the first two weeks of culture. Hepatobiliary organoids retain hepatocyte functions during expansion and recapitulate key aspects of liver tissue architecture, including the arrangement of hepatocytes into cord-like structures which terminate in biliary-like structures. As a proof-of-concept, we show that HBOs can recapitulate hallmarks of cholestasis and steatosis *in vitro*. Finally, we demonstrate that hepatocytes can be transfected, transduced and gene edited in 3D prior to organoid generation, facilitating a wide range of applications.

## Results

### Mass Generation of Hepatobiliary Organoids from Adult Primary Human Hepatocytes

We tested a panel of growth factors, inflammatory cytokines and small molecules (Table S1) targeting pathways involved in liver regeneration for their ability to stimulate the expansion of single adult PHH in 2D, 2.5D and 3D. These methods showed limited success, with 2D cultured hepatocytes proliferating minimally and displaying the prototypic signs of dedifferentiation. Hepatocytes cultured in 2.5D and 3D retained their morphology for weeks, but failed to establish expandable cultures. In some conditions we noted the aggregation of hepatocytes into clusters, which appeared more proliferative and robust compared to single hepatocytes. However, under the media conditions tested, clusters failed to expand beyond a diameter of 50-100 µm. To further probe the impact of aggregation on hepatocyte proliferation, we employed an ultra-low attachment (ULA) microwell system to promote the formation of hepatocyte spheroids, prior to embedding cells in Matrigel. Notably, prior aggregation of hepatocytes promoted their expansion and morphogenesis upon transfer to Matrigel (Figure 1A,B).

**Figure 1.**
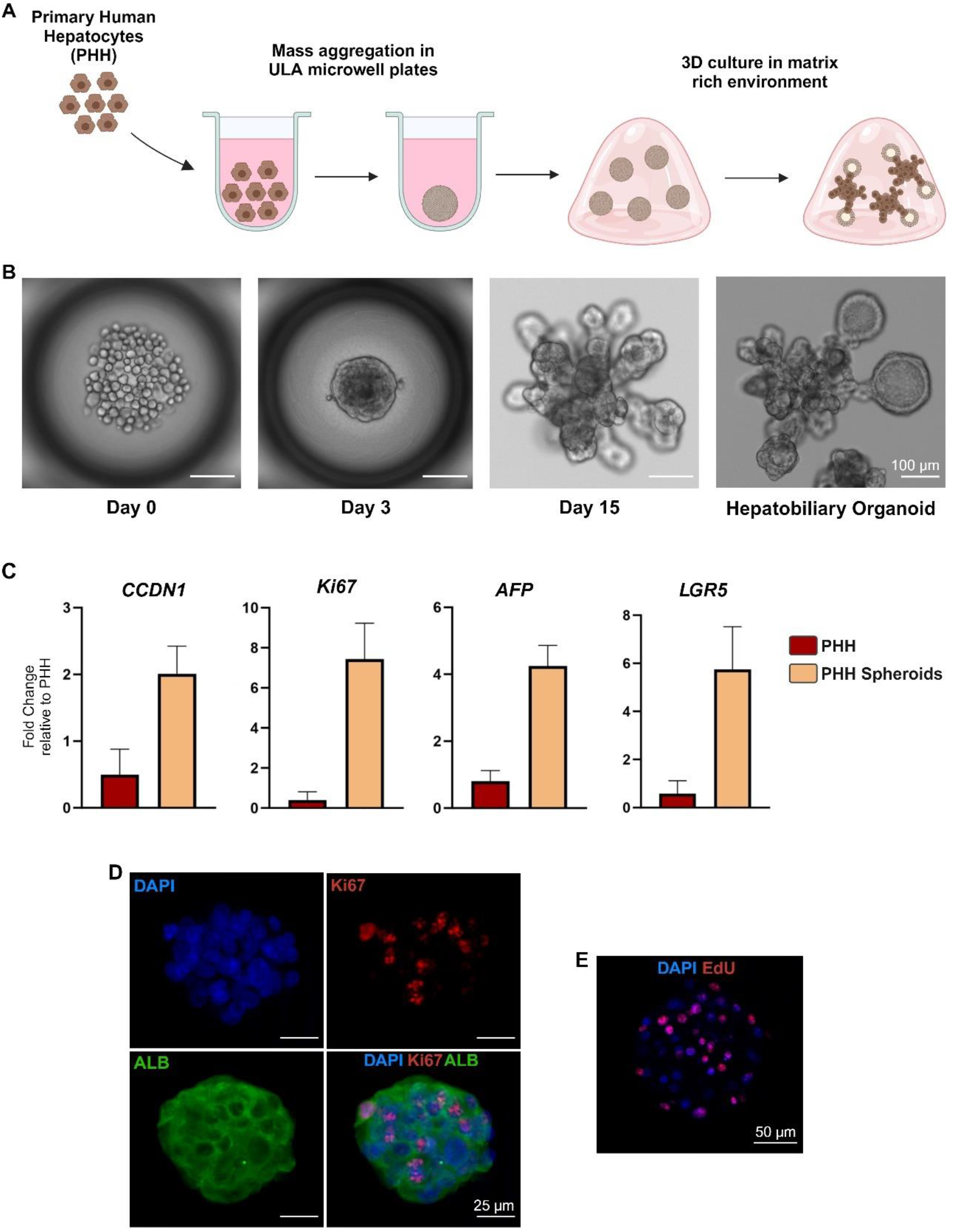
Generation of Hepatobiliary Organoids from adult PHH. A) Schematic depiction of the generation of hepatobiliary organoids from adult PHH. B) Brightfield images of PHH on day of seeding, after the formation of spheroids, and reorganization into hepatobiliary organoids (left to right), scale bars = 100 µm. C) RT-qPCR analysis of proliferation and regeneration markers in hepatocyte spheroids after 72 hours in HSM. Gene expression normalized to the mean of housekeeping genes (HMBS and RPL13a) and fold-change was set relative to donor matched PHH before culture. Graphs present mean results from 3 replicates from two independent donors (age 27 and 60). D) Confocal z-stack of hepatocyte spheroids on day 3. ALB (green), Ki67 (red), scale bar = 25 µm. E) Confocal single-plane image of hepatocyte spheroids day 3. EdU (red), scale bar = 50 µm.

After optimization, media conditions were refined to include a defined: (1) hepatocyte spheroid medium (HSM) for aggregation of hepatocytes into spheroids, (2) hepatocyte expansion medium (HEM) for further expansion of hepatocytes as hepatobiliary organoids (HBOs) and (3) hepatocyte maturation medium (HMM) to promote high functionality of HBOs.

To establish cultures adult PHH were seeded in ULA at a density of 50-75 cells per microwell and allowed to aggregate in the presence of EGF, HGF, Wnt, RSPO3, Y-27632, A83-01 and FBS (HSM medium). Under these conditions, adult PHH formed stable spheroids within 48-72 hours. The resulting hepatocyte spheroids were round and compact (Figure 1A,B). Quantitative real time PCR (RT-qPCR) revealed a strong upregulation of proliferation markers *Ki67* and *CCND1*, as well as the fetal/regeneration markers *AFP* and *LGR5* in hepatocyte spheroids compared to PHH before culture (Figure 1C). Analysis by immunofluorescence (IF) confirmed the co-expression of Ki67 and ALB on a protein level (Figure 1D), supporting the notion that hepatocytes were actively proliferating. To verify this, we performed EdU incorporation assays on hepatocyte spheroids. EdU staining showed labeling of numerous hepatocyte nuclei, confirming that the present culture conditions supported the proliferation of adult PHH (Figure 1E).

Following their formation, hepatocyte spheroids were collected and embedded in Matrigel. Over the next 2 weeks hepatocyte spheroids underwent dramatic morphological changes as hepatocytes began to interact with and expand into the matrix-rich 3D environment. In the presence of Nicotinamide, EGF, HGF, Amphiregulin, Wnt, RSPO-3, Y-27632, A83-01 and FBS (HEM medium), hepatocyte spheroids reorganized into extended, cord-like projections of hepatocytes emanating from a central region. In many organoids, hepatocyte cords terminated in clearly visible biliary-like structures (Figure 1B). H&E staining of hepatocyte spheroids and hepatobiliary organoids showed the stark organizational differences between the two (Figure 2A,B). Of note, PHH could be derived from neonatal (age 0-1) to elderly donors (age 65+) and could be sourced from commercially available vials of cryopreserved PHHs. HBOs could be expanded at a split ratio of 1:2-1:3 every 7-14 days for 2-3 months, depending on the donor (Table S2). To mature hepatobiliary organoids for functional analysis and *in vitro* modeling, growth factors (EGF, Amphiregulin, Wnt, RSPO-3), Nicotinamide and FBS were withdrawn from the medium and dexamethasone was added (HMM medium).

**Figure 2.**
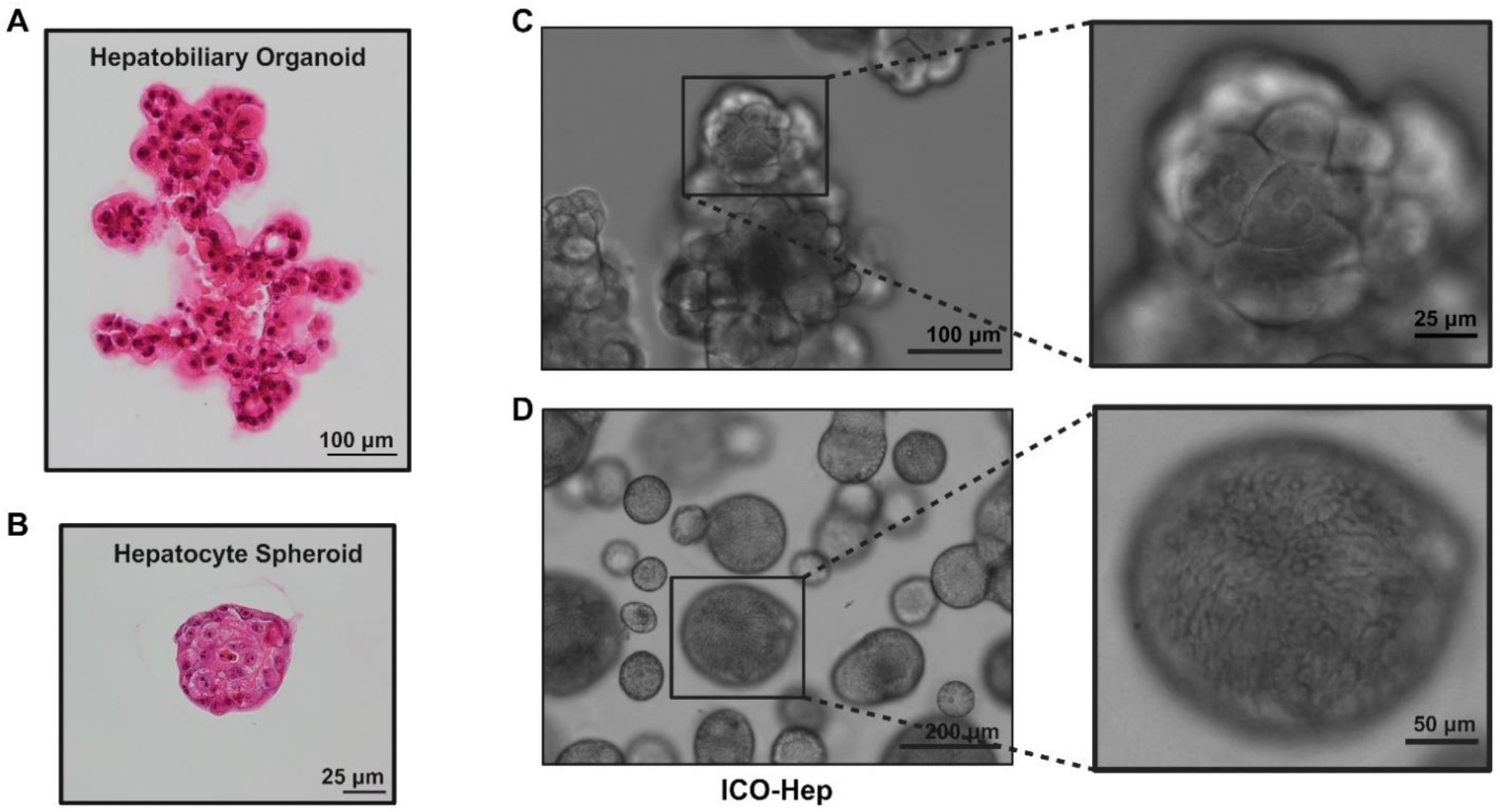
Organizational and Morphological Differences Between Hepatocyte Spheroids, HBOs and ICO-Heps. A) H&E of hepatobiliary organoid, scale bar = 100 µm B) H&E of hepatocyte spheroid, scale bar = 25 µm C) Brightfield images of HBOs on day 24 of culture in HEM (60-year-old donor). Lower magnification (left), scale bar = 100 µm. Higher magnification (right), scale bar = 25 µm. D) Brightfield images of ICO-Heps on day 13 of differentiation. Lower magnification (left), scale bar = 200 µm. Higher magnification (right), scale bar = 50 µm.

### Characterization of Hepatobiliary Organoids

To compare our HBOs to previously published organoid systems derived from primary tissue of the adult human liver, we generated donor-matched hepatocyte organoids (HOs) and intrahepatic cholangiocyte organoids (ICOs) using previously established protocols^19,24^. HOs established from single adult hepatocytes failed to expand beyond a diameter of 50-100 µm or undergo hepatobiliary morphogenesis (Figure S2A). As previously reported, <1% of adult PHH gave rise to organoids with limited expansion potential^19^, hampering in-depth analysis of the resulting colonies. In contrast, ICOs established from single adult EpCAM^+^ liver cells were capable of long-term expansion as 3D cystic structures (Figure S2C), which could be differentiated towards hepatocyte-like cells (ICO-Hep). This permitted us to perform a side-by-side comparison between hepatobiliary organoids and ICO-Heps generated from the same donor.

The cells comprising hepatobiliary organoids were of typical hepatocyte morphology, with numerous binucleated cells visible (Figure 2C). In contrast, ICOs differentiated into hepatocyte-like cells failed to acquire a clear hepatocyte morphology (Figure 2D). Immunofluorescent analysis of HBOs showed broad expression of hepatocyte markers ALB and ECAD (Figure 3A and Video S1), HNF4A (Figure 3B), as well as ASGR1 (Figure 3C). The canalicular transporter, MRP2, lined regions between hepatocytes, as well as the luminal membrane of the biliary structures (Figure 3D). EdU incorporation assays confirmed that HBOs were actively proliferating in HEM (Figure 3E). HBOs also contained cells positive for the cholangiocyte marker KRT19. Interestingly, KRT19 cells were not of typical cholangiocyte size and morphology. Instead, KRT19 cells appeared to be large, binucleated cells, which co-expressed hepatocyte markers, ALB and HNF4A (Figure 4A,B and Video S2). We also noted the emergence of cells reminiscent of the ‘small hepatocytes’ described by Mikita and colleagues in cultures of primary rat hepatocytes^25^ after long-term culture of HBOs in HEM (Figure 4C). Of note, when matured in HMM, the cells comprising the biliary structures of HBOs resembled large, mature hepatocytes (Figure 4D).

**Figure 3.**
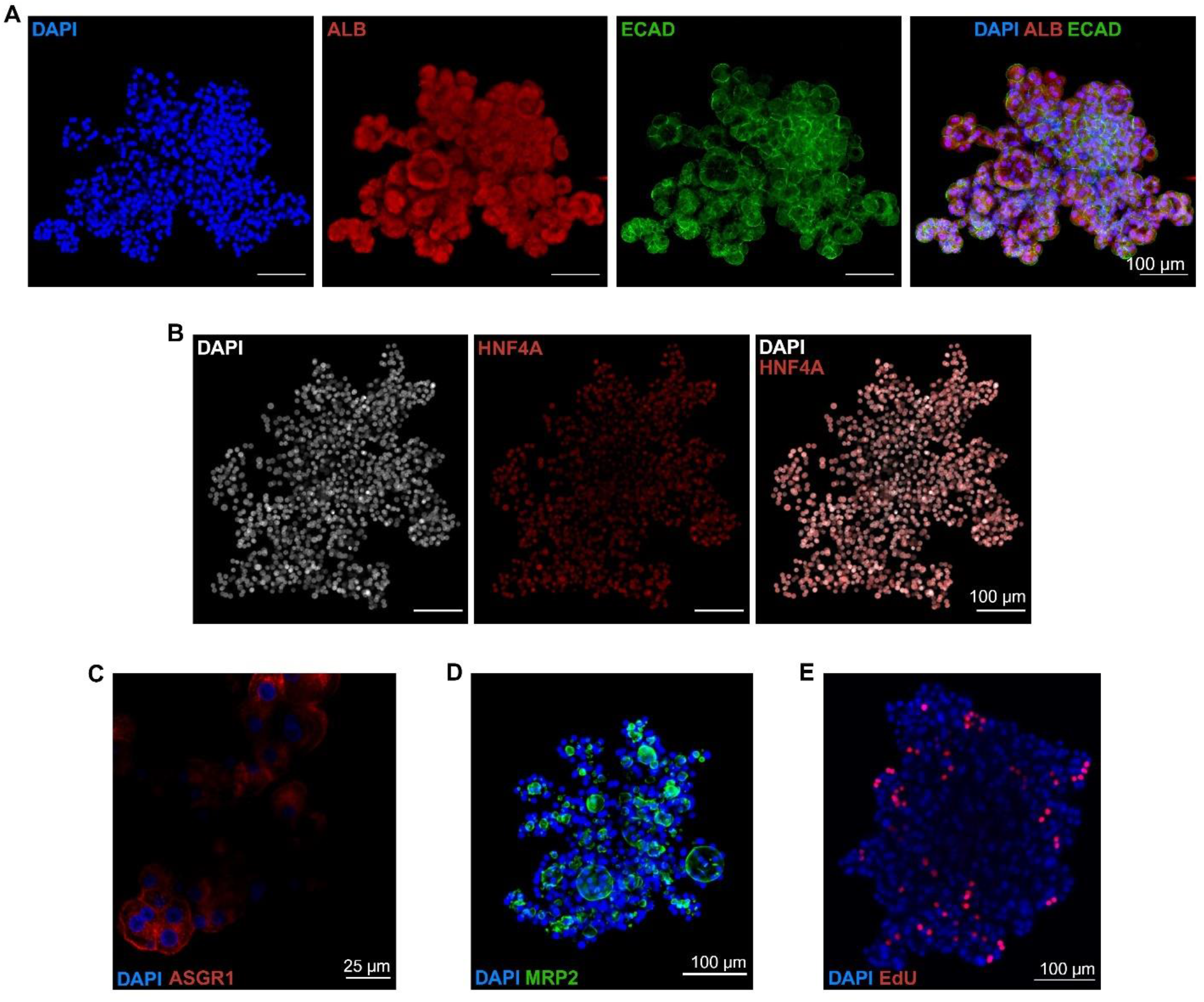
Immunofluorescent Analysis of Hepatocyte Markers in Hepatobiliary Organoids. A) Confocal z-stack of hepatocyte markers in hepatobiliary organoids in HEM. ALB (red), ECAD (green), scale bar = 100 µm. B) Confocal z-stack of hepatocyte marker HNF4A (red) in HBOs in HEM, scale bar = 100 µm. C) Confocal single-plane image of hepatocyte marker ASGR1 (red) in HBOs in HEM, scale bar = 25 µm. D) Confocal z-stack of hepatocyte marker MRP2 (green) in HBOs in HEM, scale bar= 100 µm (right). E) Confocal z-stack of hepatobiliary organoids incubated with EdU (red) on day 45 of culture in HEM, scale bar = 100 µm.

**Figure 4.**
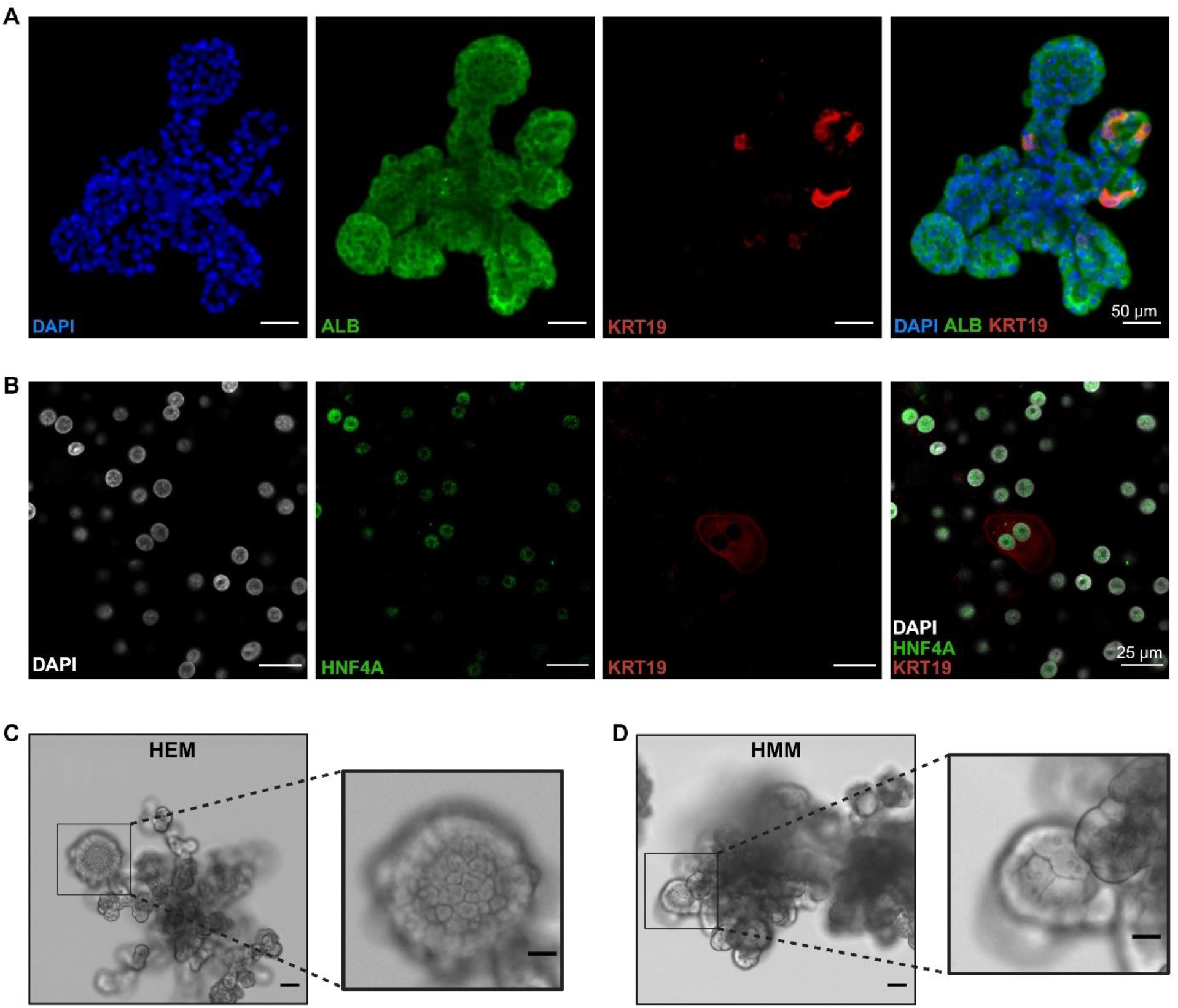
Characterization of Bi-phenotypic Cells in Hepatobiliary Organoids. A) Confocal z-stack of hepatocyte (ALB, red) and cholangiocyte (KRT19, green) markers in HBOs in HEM, scale bar = 100 µm. B) Single-plane image of hepatocyte (HNF4A, green) and cholangiocyte (KRT19, red) markers in HBOs in HEM, scale bar = 25 µm C) Brightfield images of hepatobiliary organoids on day 21 of culture in HEM. Lower magnification (left), scale bar = 50 µm. Higher magnification (right), scale bar = 25 µm. D) Brightfield images of hepatobiliary organoids on day 21 of culture in HMM. Lower magnification (left), scale bar = 50 µm. Higher magnification (right), scale bar = 25 µm.

Gene expression analysis by RT-qPCR revealed that expanding hepatobiliary organoids maintained expression of hepatocyte markers (*HNF4A, ALB, ASGR1)* (Figure 5A), although lower than before culture, while also acquiring the expression of proliferation and regeneration markers (*Ki67* and *LGR5*) (Figure 5B), as well as the biliary marker, *KRT7* (Figure 5C). HBOs in HEM also expressed key drug metabolizing genes (*CYP1A2, CYP3A4, CYP2B6, CYP2D6, CYP2E1* and *CYP2C9*) (Figure 5D) and canalicular transporters (*MRP2* and *BSEP*) (Figure 5E). Of note, the expression of hepatocyte genes in ICOs and ICO-Heps was significantly lower compared to HBOs, while the expression of biliary genes was significantly higher. When HBOs were switched from expansion (HEM) to maturation (HMM) media, expression of hepatic functional genes increased to levels comparable to PHH, while the expression of proliferation and biliary markers decreased (Figure 5). Taken together, expansion of HBOs in HEM promoted a regeneration-associated gene expression profile, which was accompanied by the partial de-differentiation of a subset of hepatocytes into cells with a hybrid hepato-biliary phenotype. Maturation of hepatobiliary organoids in HMM restored expression of hepatic functional genes and decreased expression of regeneration and biliary genes, closely resembling PHH before culture.

**Figure 5.**
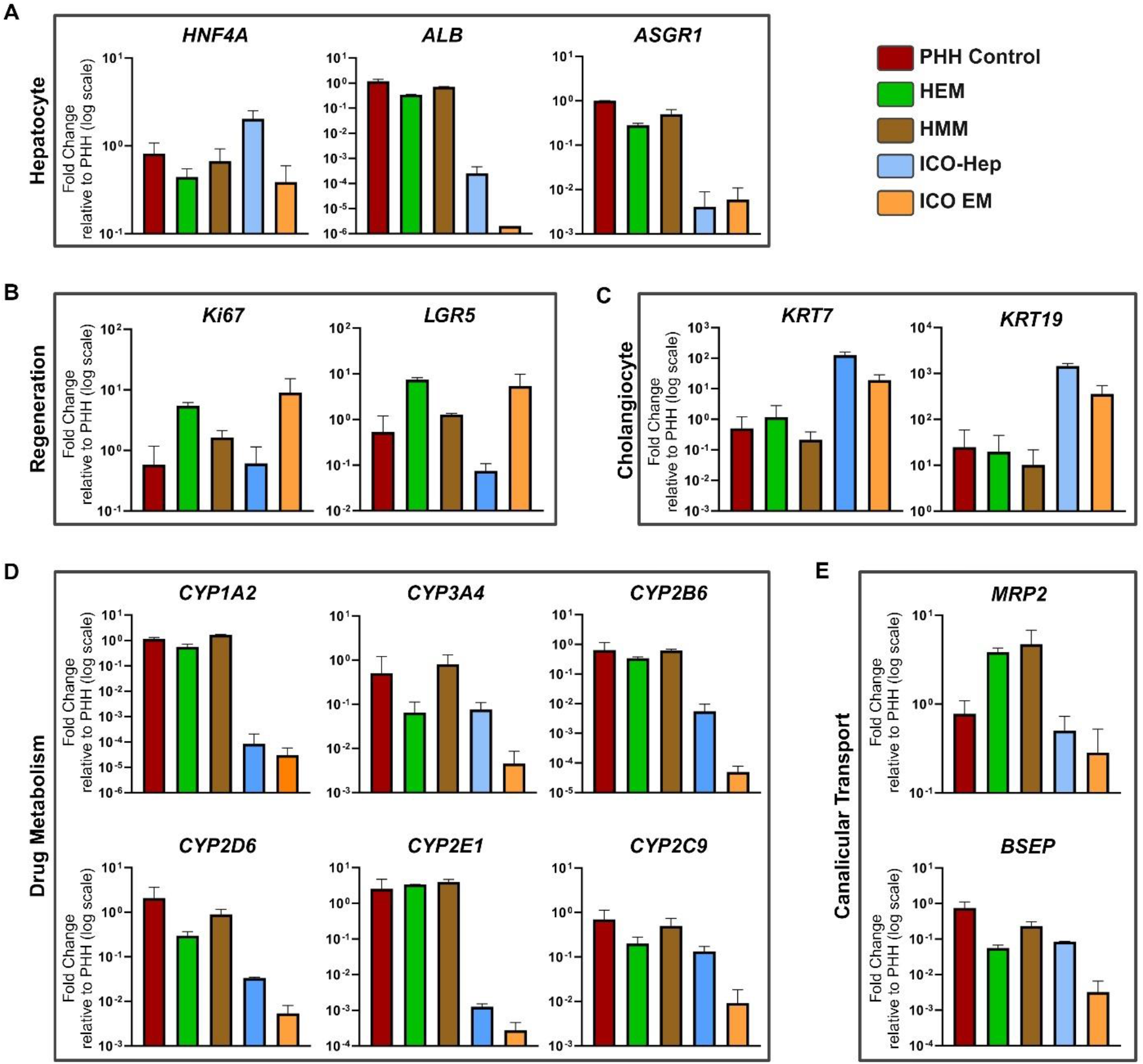
Expression of Hepatocyte Lineage, Functional, Regeneration and Biliary markers in HBOs. A) RT-qPCR analysis of hepatocyte markers. B) RT-qPCR analysis of cholangiocyte markers. C) RT-qPCR analysis of liver regeneration markers. D) RT-qPCR analysis of phase I and II drug metabolizing enzymes. E) RT-qPCR analysis of bile acid transporters. All graphs represent donor-matched PHH (control), HBOs (day 20) (HEM/HMM), ICOs (p4) and ICO-Heps (p4). Graphs present mean results from 3 replicates from two independent donors (age 27 and 60). Gene expression normalized to the mean of housekeeping genes (HMBS and RPL13A) and fold-change was set relative to donor-matched PHH before culture.

### Hepatic Functionality of Hepatobiliary Organoids

Hepatobiliary organoids were able to perform key hepatocyte functions while expanding. These include (1) glycogen storage, demonstrated by periodic acid-Schiff (PAS) staining (Figure 6A), (2) uptake of fluorescently labeled low-density lipoproteins (Figure 6B), (3) albumin secretion, (4) canalicular transport and (5) metabolism of xenobiotics through phase I and II drug metabolizing enzymes. Consistent with gene expression analysis, HBOs secreted more albumin following maturation in HMM (Figure S3). When compared to donor matched ICO-Heps, hepatobiliary organoids in HMM secreted at least 1000 times more albumin, with levels comparable to 2D PHH in sandwich culture (Figure 6C). To visualize canalicular transport, HBOs were incubated with 5-(and-6)-carboxy-2′,7′-dichlorofluorescein diacetate (CDFDA), which is cleaved by intracellular esterases to yield the MRP2 substrate, 5 (and 6)-carboxy-2′,7′-dichlorofluorescein (CDF)^26^. Hepatobiliary organoids were able to readily transport CDF, which accumulated in canalicular regions within the organoids, as well as the lumen of the interconnected biliary-like structures (Figure 6D). To determine drug metabolizing capabilities, hepatobiliary organoids were treated with a cocktail of substrates of phase I (CYP3A4, CYP2C9, CYP2B6, CYP2D6, CYP1A2) and phase II (UGTs) enzymes. Parent compound depletion and metabolite formation were measured by LC-MS/MS. This revealed successful formation of each metabolite in both expanding (HEM) and matured (HMM) HBOs (Figure 6E). Of note, the activity of all CYP enzymes assayed increased following maturation of HBOs. For example, acetaminophen production from CYP1A2 was up to 20 times higher in hepatobiliary organoids following maturation. Furthermore, CYP activity was significantly higher in HBOs than in ICO-Heps, particularly the production of acetaminophen from phenacetin, which was up to 1000 times higher in matured HBOs. Taken together, hepatobiliary organoids retain key hepatocyte functions while actively proliferating. Induction of maturation through the withdrawal of proliferative cues increases hepatic functionality of hepatobiliary organoids.

**Figure 6.**
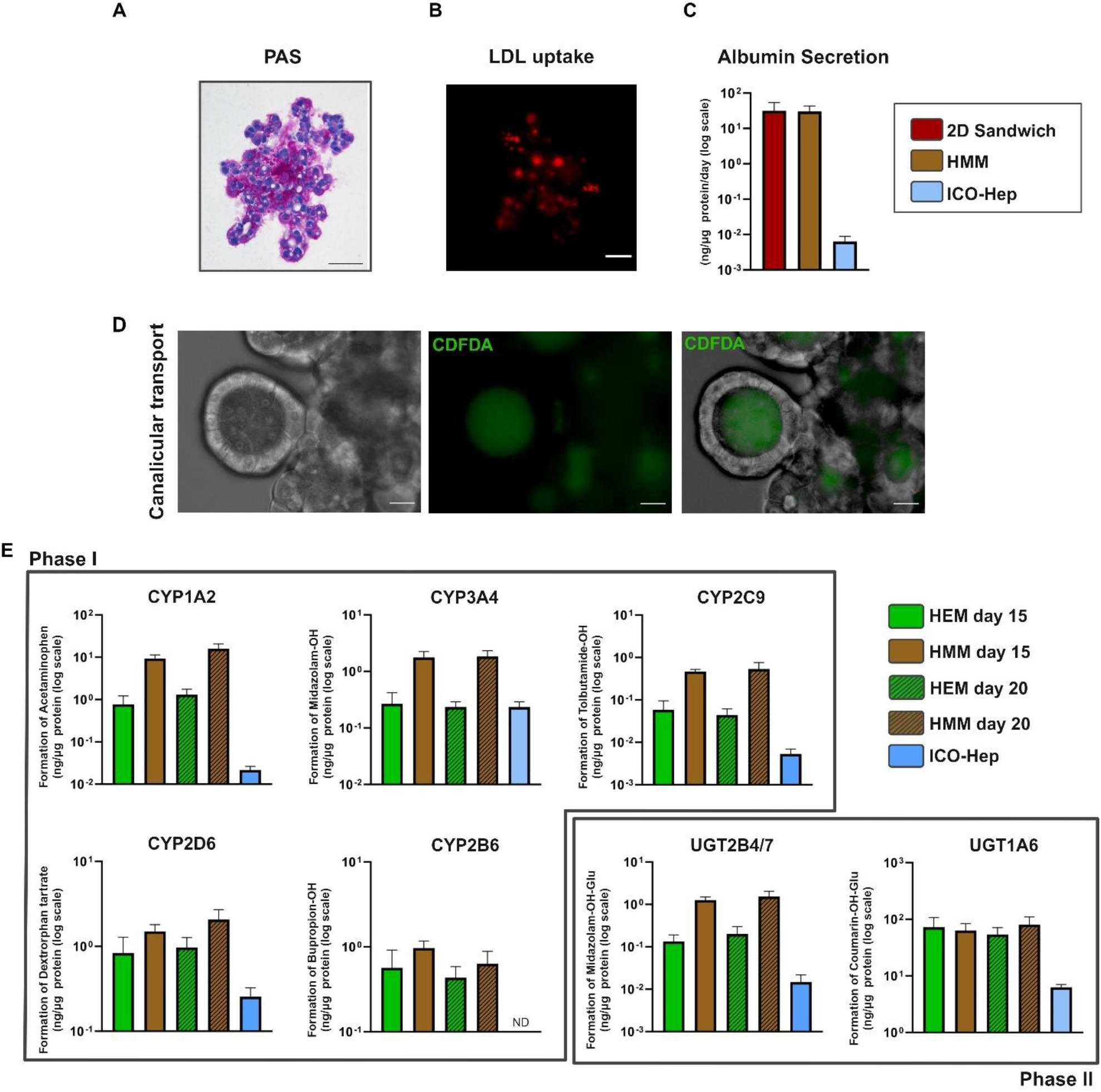
Hepatic Functionality of Hepatobiliary Organoids. A) Glycogen accumulation in HBOs in HEM detected by Periodic-Acid Schiff (PAS) staining, scale bar = 100 µm. B) Low density lipoprotein uptake in HBOs in HEM visualized with Image-iT™ Low Density Lipoprotein Uptake Kit, pHrodo™ (red), scale bar = 100 µm. C) Albumin secretion measured after 24 hours (2D sandwich culture, day 2 of culture) or 48 hours in HBOs (Day 20 of culture, HMM), ICO-Heps (p4, DM day 13). Graph presents mean results of 3 replicates from 3 individual wells of a single donor. Results are presented as ng albumin/µg protein/day. D) Canalicular transport of CDFDA [1µM]. Brightfield (left), CDFDA (middle), merge (left), scale bar = 25 µm. E) Formation of metabolites from phase I (top) and II (bottom) enzymes after 48 hours in HBOs (Day 15 and 20 of culture (HEM and HMM) and ICO-Hep (p4, DM day 13). Graph presents mean results of 3 replicates from 3 individual wells of a single donor. Results are presented as ng/µg protein.

### Recapitulating Hallmarks of Cholestasis and Steatosis in Hepatobiliary Organoids

To probe the utility of hepatobiliary organoids as a model for cholestasis, we set out to determine whether organoids could uptake and transport bile acids. Multiple luminal structures could be seen at the end of hepatocyte cords, reminiscent of canals of Hering (Figure 7A). Immunofluorescent analysis revealed that the bile acid transporter, BSEP, localized to canalicular regions between hepatocytes (Figure 7B). To visualize transport of bile acids HBOs were incubated with the fluorescent bile acid analogue, Cholyl-L-lysyl-fluorescein (CLF) [1 µM], which is widely used to visualize canalicular transport of bile acids within liver tissue. Hepatobiliary organoids were able to readily uptake and transport CLF, which accumulated in canalicular regions within the organoids (Figure 7C). Of note, treatment with the cholestasis inducing drug, troglitazone (TGZ), markedly reduced canicular accumulation of CLF (Figure 7C), demonstrating that bile acid transport could be effectively disrupted.

**Figure 7.**
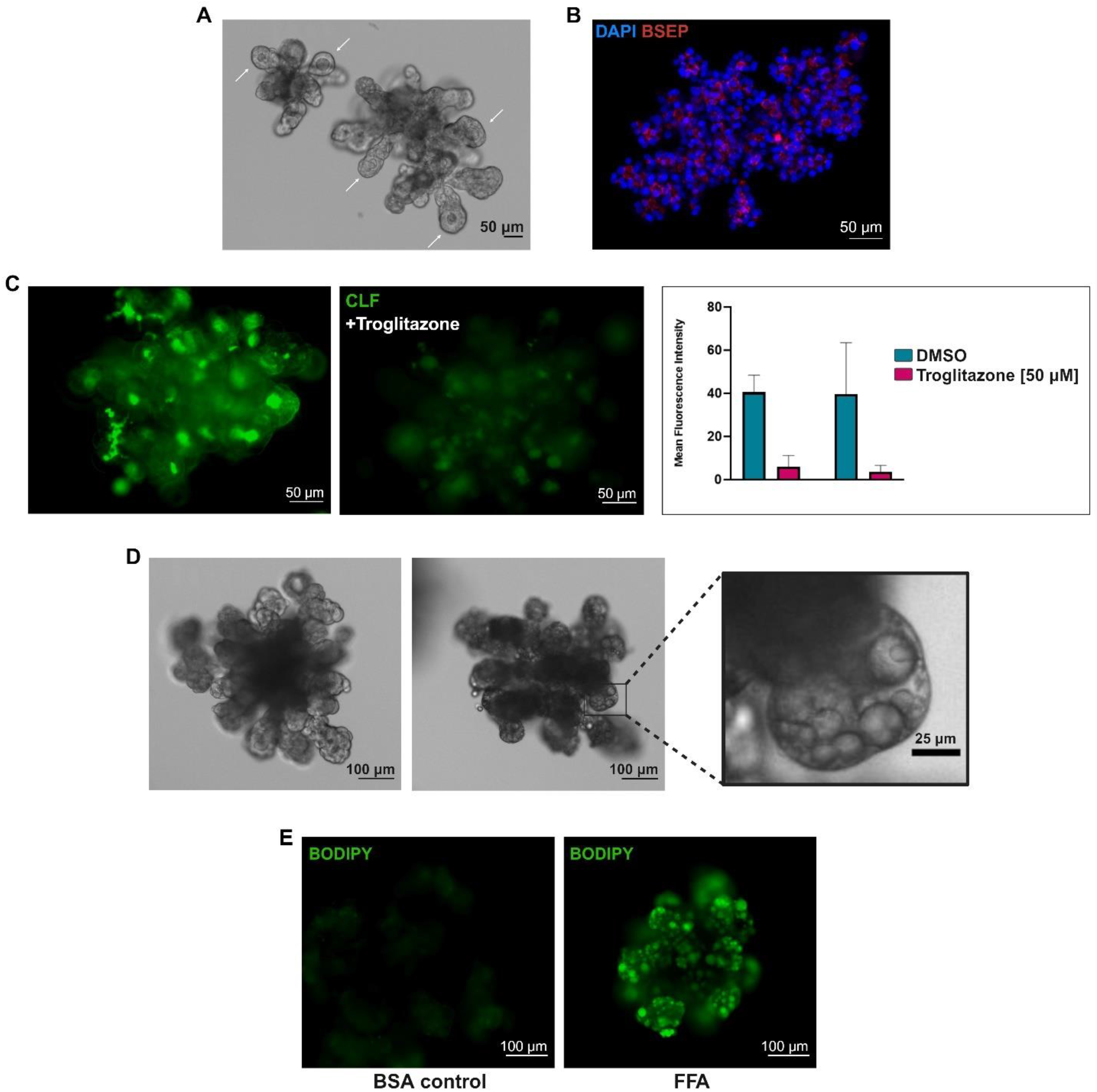
Modeling Cholestasis and Steatosis. A) Brightfield image of hepatobiliary Organoids in HEM (day 10), scale bar = 50 µm. White arrows indicate luminal structures at the end of hepatocyte cord-like structures. B) Confocal z-stack of bile acid transporter (BSEP, red) in hepatobiliary organoids, scale bar = 100 µm. C) Single-plane image of hepatobiliary organoids incubated with CLF [1 µM] (left) or CLF [1 µM] + troglitazone [50 µM] (right). CLF (green), scale bar 50 µm. Graph presents mean fluorescence intensity from 10 organoids in each condition. D) (Top) Brightfield images of hepatobiliary organoids on day 12 of culture in HMM + BSA (left) or lipid cocktail (right). Lower magnification scale bar = 100 µm. Higher magnification scale bar = 25 µm. E) Single-plane image of live hepatobiliary organoids on day 12 of culture in HMM + BSA (left) or lipid cocktail (right) stained with BODIPY 493/503 [1 µg/mL], scale bar = 100 µm.

Another pathological liver condition, steatosis, is characterized by the accumulation of lipid droplets (LDs) in the cytoplasm of hepatocytes. To determine whether HBOs could be used to model steatosis we matured organoids in HMM for 4 days and then exposed them to a cocktail of lipids for 12 days. Hepatobiliary organoids accumulated lipids over time, generating LDs of various sizes over the time course of treatment (Figure 7D,E). Live cell staining of HBOs with the lipophilic fluorescent dye, BODIPY, showed the massive accumulation of lipids in the treatment group compared to the BSA control (Figure 7E). Taken together, we have provided proof-of-principle that HBOs can recapitulate fundamental hallmarks of cholestasis and steatosis, including bile acid transport disruption and lipid droplet accumulation, respectively.

### Transduction, Gene editing and Transfection of Hepatocytes in 3D

The ability to edit the genome of adult PHH *in vitro* could facilitate the establishment of mutation specific disease models, such as monogenetic liver diseases, as well as the correction of patient-derived cells for use as ATMPs. It was previously reported that PHHs could be infected with recombinant adenovirus prior to aggregation and spheroid formation^12^. To determine whether a similar approach could also be used to edit the genome of adult PHHs prior to the generation of hepatobiliary organoids, we seeded adult PHH in ULA microwells in HSM containing engineered virus-like particles (eVLPs) carrying a base editor targeting Heparin Binding EGF Like Growth Factor (HB-EGF). Transduction of gene editing machinery during aggregation resulted in bulk editing efficiency up to 68% (Figure 8A). Importantly, edited hepatocyte spheroids were viable and capable of generating hepatobiliary organoids following transfer to Matrigel (Figure 8B).

**Figure 8.**
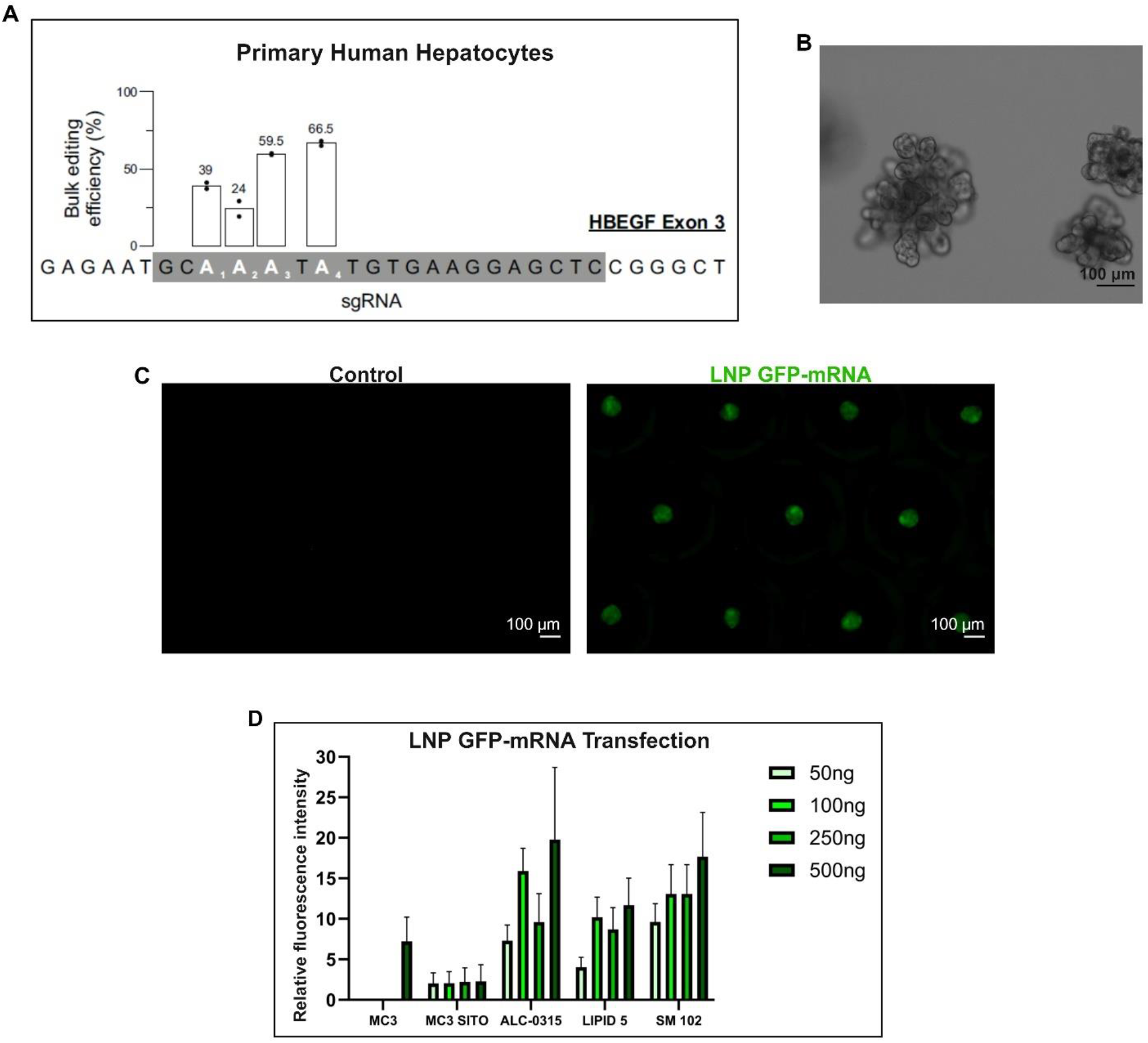
Transduction, Gene Editing and Transfection of Hepatocytes. A) Bulk editing efficiency of PHH transduced with eVLPs carrying base editor targeting HBEGF exon 3. Graph presents mean results of 2 individual wells of a single donor. B) Brightfield image of hepatobiliary organoid generated from gene edited (eVLPs, HBEGF) hepatocytes, scale bar = 100 µm. C) Single-plane images of hepatocyte spheroids at 72 hours (control, left) following transfection with LNP (ALC-0315) (100ng) carrying GFP-mRNA (right). GFP (green), scale bar = 100 µm. E) Fluorescence intensity after transfection five LNP formulations (MC3, MC3-Sito, ALC-0315, Lipid 5 and SM102) at 4 doses (50, 100, 250 and 500 ng). Graph presents mean fluorescence intensity from 15 organoids in each condition.

Next, we set out to determine whether the methodology could also be extended to other cell delivery strategies, such as transfection with lipid nanoparticles (LNPs), a technique in clinical trials for *in vivo* mRNA delivery to treat propionic acidaemia^27^ and methylmalonic acidemia^28^. In order to do this, we treated hepatocytes with five lipid nanoparticle (LNP) formulations encapsulating GFP-mRNA during aggregation in HSM. After 72 hours the resulting hepatocyte spheroids were imaged (Figure 8C), and mean GFP fluorescence intensity was determined. Transfection of PHH with LNPs carrying GFP-mRNA resulted in a dose dependent increase in fluorescence intensity, which varied depending on the LNP formulation (Figure 8D). Taken together, we demonstrate that adult PHH can be efficiently gene edited and transfected *in vitro* in 3D, facilitating a wide range of applications, such as the establishment of monogenetic liver disease models and the optimization of LNP formulations targeting the liver.

## Discussion

Adult primary human hepatocytes are the cell source of choice for hepatic toxicity testing, disease modeling and regenerative medicine applications. While the differentiation of pluripotent stem cells (PSCs) into hepatocytes has seen marked improvement over recent years, no protocol to date has been able to generate cells comparable to adult PHH, specifically with regard to functionality. However, to truly exploit the potential of PHH, novel methods to maintain and expand these cells *in vitro*, as well as generate faithful and predictive 3D tissue-like models, are required. Recent efforts have focused on (1) single-cell expansion of PHHs as epithelial organoids in a matrix-rich 3D environment^19,20,29^ and (2) aggregation of hepatocytes into spheroids in an environment devoid of exogenously supplemented extracellular matrix (ECM) cues^12,22,23^. The former approach has succeeded in establishing expandable organoids from fetal liver cells, but has shown limited success in the efficient generation of proliferative organoids from adult PHHs^30^. The latter method has succeeded in generating 3D liver cultures which can be maintained for 1-2 months, but has failed to recapitulate key aspects of liver tissue architecture, such as the arrangement of hepatocytes in cords which terminate in bile ducts^12^. Recently, Ramli *et. al* reported the generation of organoids from PSCs with functionally interconnected hepatic and biliary domains, however the methodology employed necessitates a lengthy differentiation protocol spanning 2-3 months for organoid generation^31^. While this work was in progress Hendricks *et al*. reported the improved growth of PHHs from single cells utilizing IL-6 and an FXR agonist. Using their methodology neonatal and infant (0.3 and 1.7 years, respectively) derived hepatocytes could be expanded for up to 4 months as cystic-like structures^20^. Interestingly, the constituent cells are reminiscent of the ‘small hepatocytes’ we also observed in the biliary structures of the currently reported hepatobiliary organoids (in expansion conditions) (Figure 4C), and which upon dissociation into single cells (followed by secondary culture in expanding conditions), reformed organoids similar to those reported by Hendricks *et al*. (Figure S4).

Consistent with previous reports^19,21,22^ and the broader literature on liver regeneration^32–35^, the activation of Wnt signaling combined with growth factor stimulation promoted the expansion of PHH. Stimulation of hepatocytes with inflammatory cytokines (IL-6 and TNFα) was not necessary for the generation of proliferative organoids, contrary to what has been described for mouse and human hepatocyte organoids derived from single cells^20,36^. During optimization of the expansion media, we tested multiple concentrations of TNFα and IL-6, individually, and in combination. In general, TNFα appeared to have negative effects on cultures, while IL-6 stimulated the expansion and regeneration of organoids, though this was accompanied by a morphological shift towards cystic-like organoids, which was further exacerbated by inclusion of the FXR agonist, cilofexor, as recently reported^20^. In general, proliferation of HBOs tended to slow after 4-6 weeks of expansion. As such, while the current effort has made progress towards the efficient expansion of adult PHH through the mass generation of expanding organoids, the full exploitation of their intrinsic regenerative potential will require further efforts. Given the fundamental role of inflammatory cytokines in liver regeneration *in vivo*, further refinements of the culture conditions may promote improved expansion of adult PHH. Of note, adult PHH have been demonstrated to possess the regenerative potential to repopulate several mouse livers *in vivo* through serial hepatocyte transplantations^37^. This indicates that the inability to efficiently expand adult PHH to clinically relevant cell numbers *in vitro* is not a result of an intrinsic regenerative limitation, but rather *in vitro* culture conditions.

While there are multiple lines of evidence indicating that select subsets of hepatocytes are responsible for liver regeneration^38–40^, recent reports demonstrate that the regenerative potential of hepatocytes is not restricted to a select subset, but is an intrinsic capability of mature hepatocytes, which can transition between multiple cell states depending on metabolic and regenerative demands^41,42^. During liver regeneration in a variety of injury contexts, hepatocytes in all liver zones contribute to liver regeneration^43–45^. This is accompanied by the functional compensation of the non-proliferative hepatocytes to keep up with metabolic demands necessary to sustain life while the liver parenchyma is restored^46^. Using this rationale, aggregation of hepatocytes into functional units prior to expansion could facilitate the proliferation of hepatocytes, which are functionally assisted by hepatocytes which maintain a more mature phenotype. Interestingly, previous single-cell analysis of hepatocyte organoids derived from single cells revealed the existence of different sub-clusters of cells, some of which were enriched for proliferation/regeneration markers, while others were enriched for functional markers^19,20^.

Here we describe the mass generation and long-term expansion of organoids derived from adult primary human hepatocytes. Rather than capturing the low percentage of hepatocytes amenable to single cell expansion, we capture the broader hepatocyte population through the mass aggregation of hepatocytes into spheroids. Upon transfer to Matrigel, hepatocyte spheroids reorganize into cord-like projections of hepatocytes with interconnected biliary-like structures, in which hepatocytes do not display their prototypic canalicular polarization, but instead form cystic structures lined by canalicular transporters. Expanding hepatobiliary organoids retain key morphological, transcriptomic and functional characteristics of PHH, while also displaying regeneration-associated bi-phenotypic features. Unlike previous studies, which report the absence of cholangiocyte markers in hepatocyte organoids^19,20,36^, hepatobiliary organoids in expansion conditions upregulate the expression of *KRT7* and contain bi-phenotypic cells which co-express KRT19 and ALB. Interestingly, Gribben *et al*. recently described the emergence of similar cells *in vivo* during end-stage-liver disease, a plasticity between cholangiocytes indicative processes^42^.

Hepatobiliary organoids can be mass produced and matured for *in vitro* modeling within two weeks, and expanded at a split ratio of 1:2-1:3 for 2-3 months. Of note, PHH can be derived from neonatal (age 0-1) to elderly donors (age 65+), and can be sourced from fresh liver tissue, as well as commercially available vials of cryopreserved PHH. Because of the high formation efficiency, it is possible to isolate hepatocytes for culture and in-depth analysis from small liver biopsies, enabling personalized medicine applications. Further, the ability to edit the genome of isolated adult PHH during aggregation at high efficiency could theoretically permit the *ex vivo* gene correction of PHH in 3D, followed by autologous transplantation of gene-corrected cells; a promising approach for monogenetic liver diseases, in which it has been estimated that replacement of a small percentage (5-10%) of liver mass is sufficient to achieve improved clinical outcomes^3^.

Taken together, we describe the mass generation and long-term expansion of organoids derived from adult PHHs, which recapitulate key functional and architectural aspects of the hepatic epithelium. The rapid mass production and expandability of organoids bridges the gap between short-term functionality of primary human hepatocytes and the need for scalable, long-term organoid models of the adult liver, offering immense potential for drug testing, disease modeling, and advanced therapeutic applications. We anticipate that the reported methodology and culture conditions could be extended to enable the generation of tumor organoids from well-differentiated hepatocellular carcinoma (HCC) specimens, facilitating the investigation of patient specific therapies. Similarly, our reported gene editing strategy could enable establishment of mutation specific HCC organoids from healthy cells for investigation of anti-tumor therapies depending on the underlying mutations.

---

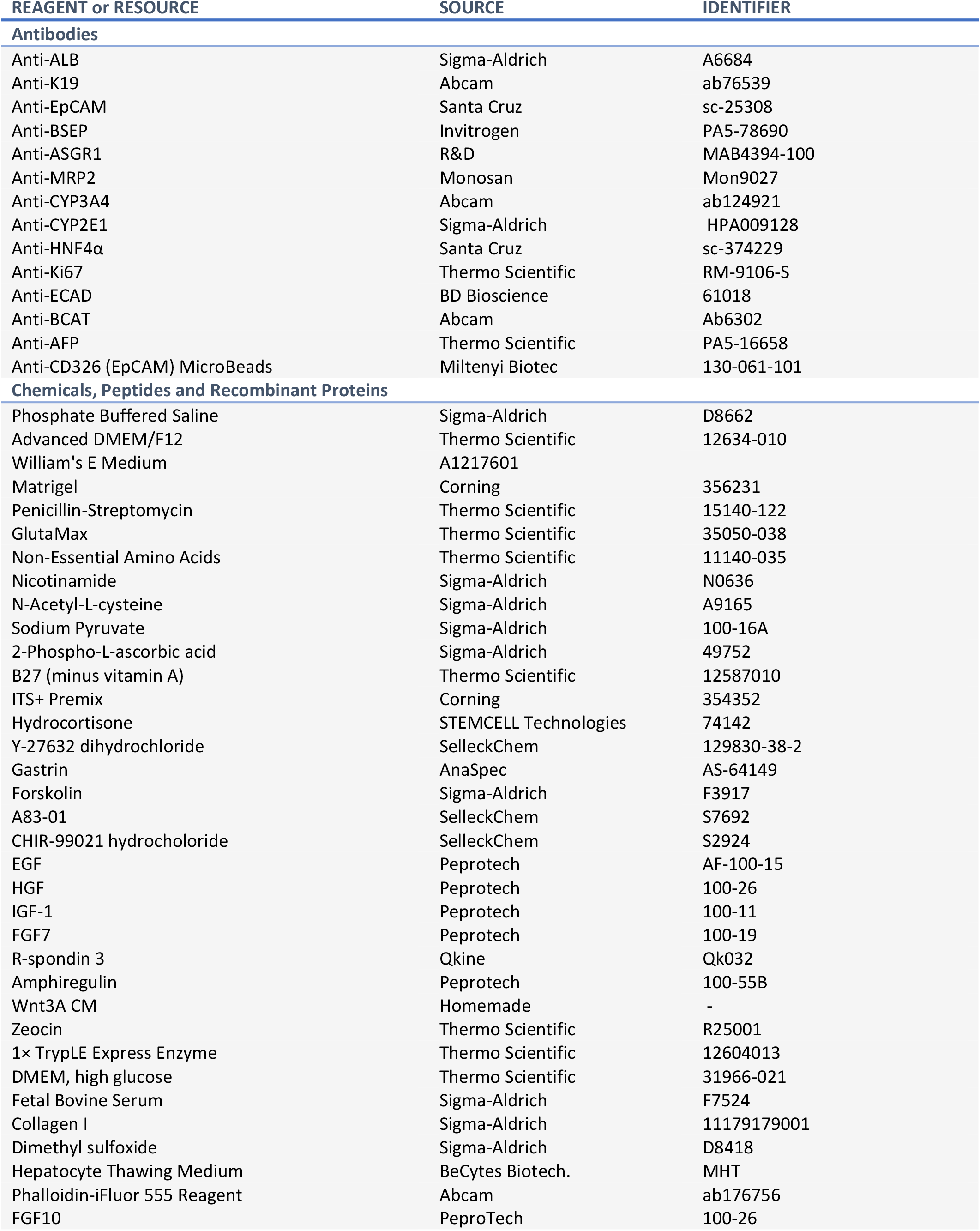
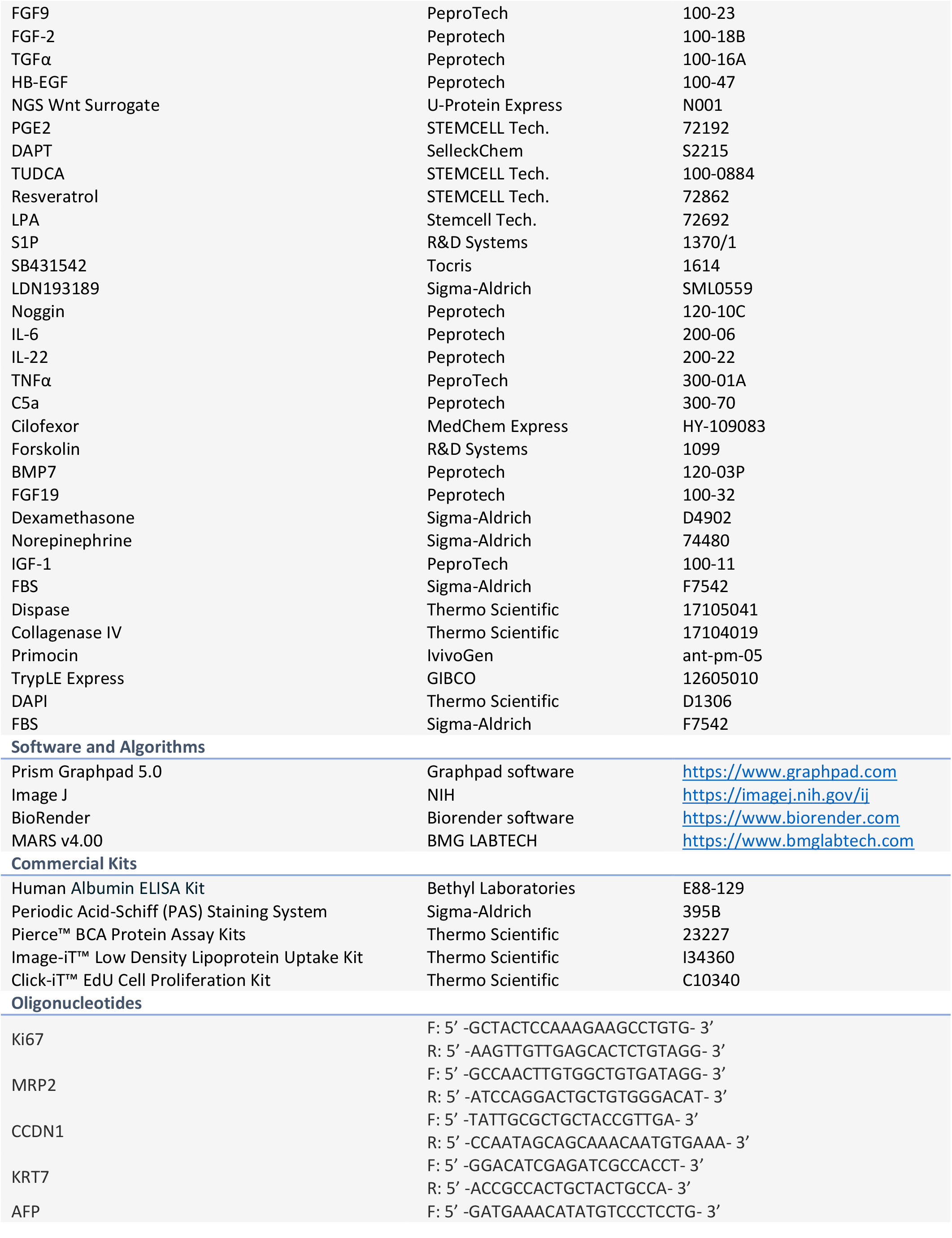
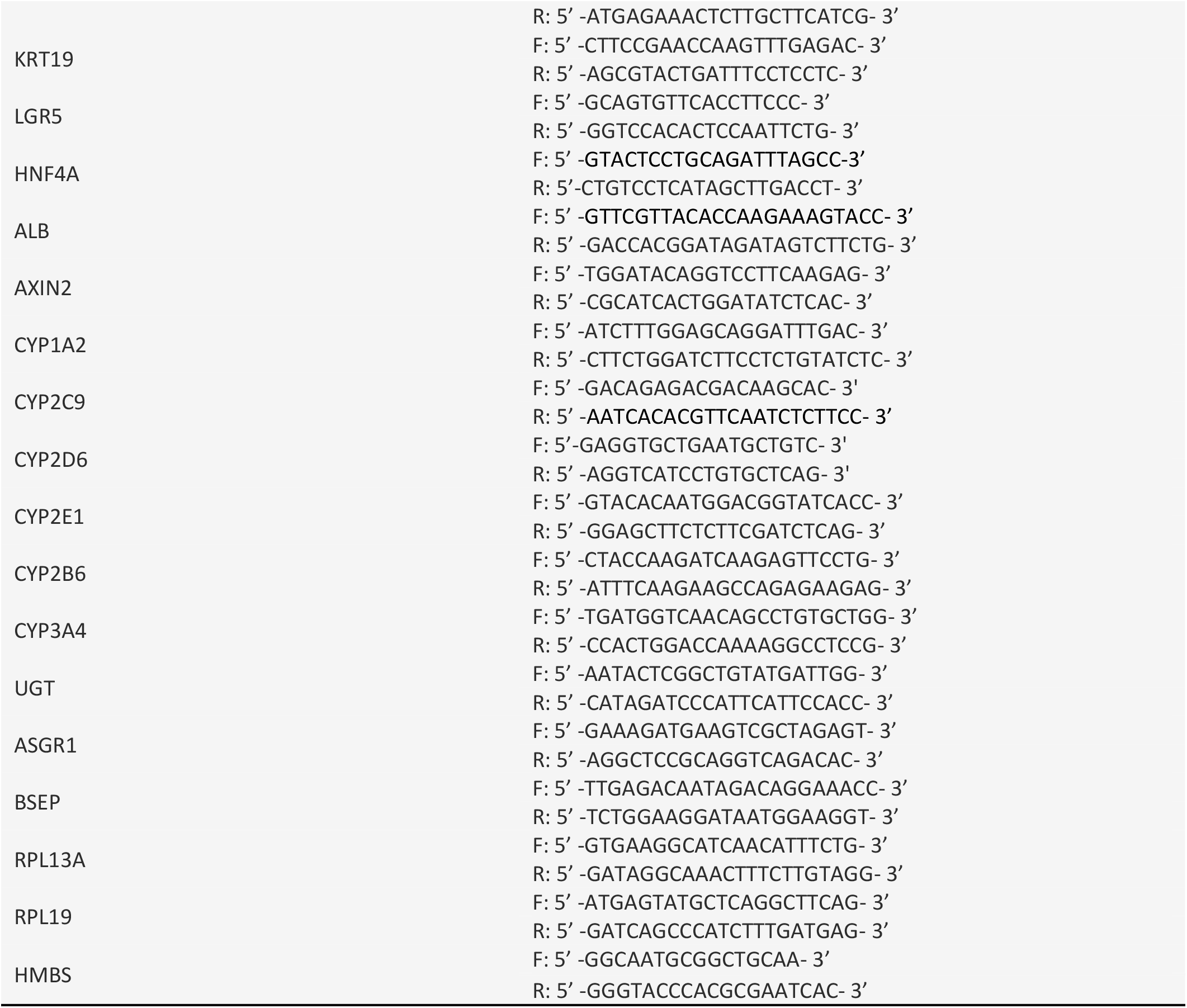

## Materials and Methods

### Isolation of primary adult human hepatocytes and cholangiocytes

Primary human hepatocytes (PHHs) and cholangiocytes (PHCs) were isolated from liver surgical resections after informed consent. Hepatocytes were isolated by two-step collagenase perfusion or mincing tissue into small fragments followed by 10-15 minutes collagenase digestion. Cell suspensions were filtered through a 70-µm cell strainer and centrifuged at 50g for 8 minutes. Cholangiocytes were isolated from small liver biopsies by mincing tissue into small fragments followed by 45-60 minutes collagenase digestion. EpCAM+ cholangiocytes were purified using EpCAM microbeads (Miltenyi Biotec) following the manufacturer’s instructions. EpCAM+ cholangiocytes were also purified from vials of cryopreserved PHH using EpCAM microbeads.

### Generation of hepatocyte spheroids

Fresh or cryopreserved primary adult human hepatocytes (BeCytes Biotechnologies, BiolVT or Sigma-Aldrich) were resuspended in hepatocyte spheroid medium (HSM). HSM consists of Advanced DMEM/F12 supplemented with 2 mM GlutaMax, 1% (v/v) Non-Essential Amino Acids Solution, 100 U/mL Penicillin-Streptomycin (all Gibco), 1 mM Sodium Pyruvate (Sigma-Aldrich), 100 µM 2-Phospho-L-ascorbic acid trisodium salt (Sigma-Aldrich), 1% (v/v) B27 (minus vitamin A) (Gibco), 1% (v/v) ITS+ Premix (Corning), 1 mM N-acetylcysteine (Sigma-Aldrich), 10 ng/mL EGF (Peprotech), 5 ng/mL HGF (Peprotech), 250 ng/mL RPSO3 (Peprotech), 0.5 nM Wnt Surrogate (IPA Therapeutics), 10 µM γ-27632 (Selleckchem), 2 µM A-83-01 (Selleckchem) and 5% (v/v) FBS (Sigma-Aldrich). Cells were plated in ultra-low attachment microwell cavities (Corning) at a density of 50-75 cells per microwell in HSM and allowed to form spheroids over 72 hours.

### Generation and expansion of hepatobiliary organoids

Hepatocyte spheroids generated in HSM were collected and centrifuged at 50g for 5 minutes. The supernatant was removed and spheroids were resuspended in 7 mg/mL Matrigel solution obtained by mixing Matrigel (Corning) with hepatocyte expansion medium (HEM) lacking FBS, B-27 and ITS+ Premix. Hepatocyte spheroids were plated at a density of 80-90 spheroids per 50 uL Matrigel droplet. Following polymerization of Matrigel to a stable hydrogel. HEM was added to each well. HEM consists of Advanced DMEM/F12, supplemented with 2 mM GlutaMax, %1 (v/v) Non-Essential Amino Acids Solution, 100 U/mL Penicillin-Streptomycin (all thermo), 1 mM Sodium Pyruvate (Sigma-Aldrich), 100 µM 2-Phospho-L-ascorbic acid trisodium salt (Sigma-Aldrich), 1% (v/v) B27 (minus vitamin A) (Thermo), 1% (v/v) ITS+ Premix (Corning), 10 mM Nicotinamide (Sigma-Aldrich), 1 mM N-acetylcysteine (Sigma-Aldrich), 10 nM gastrin (Sigma-Aldrich), 50 ng/mL EGF (Peprotech), 25 ng/mL HGF (Peprotech), 250 ng/mL RPSO3 (Peprotech), 0.5 nM Wnt Surrogate (IPA Therapeutics), 25 ng/mL Amphiregulin (Peprotech), 2 nM glucagon (ProSpec), 10 uM γ-27632 (Selleckchem), 2 uM A-83-01 (Selleckchem) and 1-5% (v/v) FBS (Sigma-Aldrich). After 1-2 weeks, Matrigel was digested with dispase to release intact hepatobiliary organoids, which were embedded in new Matrigel at a 1:2-1:3 split ratio. Organoids were passaged in this manner for ∼1 month. After which organoids were gently dissociated into small clumps of cells through a further incubation with 1x TrypLE Express (GIBCO) every other week and passaged at a 1:2-1:3 split ratio for ∼1-2 months. Medium was refreshed every 3-4 days throughout the culture period. Pro-longed enzymatic incubations, mechanical shearing, or single-cell dissociation of hepatocytes was detrimental to the successful generation of organoids.

### Maturation of hepatobiliary organoids

To mature hepatobiliary organoids, HEM was withdrawn and hepatocyte maturation media (HMM) was added. HMM consists of William’s E Medium supplemented with 2 mM GlutaMax, 1% (v/v) Non-Essential Amino Acids Solution, 100 U/mL Penicillin-Streptomycin, 1% (v/v) B27 (minus vitamin A) (all Thermo), 1% (v/v) ITS+ Premix (Corning), 1 mM N-acetylcysteine (Sigma-Aldrich), 5 ng/mL HGF (Peprotech), 2 nM glucagon (ProSpec), 10 µM γ-27632 (Selleckchem), 2 µM A-83-01 (Selleckchem) and 100 nM Dexamethasone (Sigma-Aldrich). Medium was refreshed every 3-4 days.

### 2D sandwich culture of PHH

Adult PHHs were resuspended in hepatocyte plating medium (HPM) and plated on collagen-coated plates. HPM consists of William’s E Medium supplemented with 2 mM GlutaMax, 1% (v/v) Non-Essential Amino Acids Solution, 100 U/mL Penicillin-Streptomycin (all Thermo), 1% (v/v) ITS+ Premix (Corning), 1 mM N-acetylcysteine, 100 nM Dexamethasone, and 5% (v/v) FBS (all Sigma-Aldrich). After 6-8 hours the medium was switched to HPM lacking FBS. Medium was refreshed daily.

### Generation, expansion and differentiation of intrahepatic cholangiocyte organoids

Single EpCAM+ cholangiocytes were resuspended in 7 mg/mL Matrigel solution obtained by mixing Matrigel (Corning) with cholangiocyte expansion medium (CEM) lacking B-27 and N-2. Cells were plated at a density of 500-1000 cells per 50 uL Matrigel droplet. Following polymerization of Matrigel to stable hydrogel CEM was added to each well. CEM was made as previously described (Huch., 2015): advanced DMEM/F12, supplemented with 2 mM GlutaMax, 1% (v/v) Non-Essential Amino Acids Solution, 100 U/mL Penicillin-Streptomycin, 10 mM HEPES (all Gibco), 1 mM Sodium Pyruvate (Sigma-Aldrich), 100 µM 2-Phospho-L-ascorbic acid trisodium salt (Sigma-Aldrich), 1% (v/v) B27 (minus vitamin A) (Thermo), 1% (v/v) N2 (Thermo), 10 mM Nicotinamide, 1 mM N-acetylcysteine, 10 nM gastrin (all Sigma-Aldrich), 100 ng/mL FGF10 (Peprotech), 50 ng/mL EGF (Peprotech), 25 ng/mL HGF (Peprotech), 10% RSPO1 CM (homemade), 5 µM A-83-01 (Selleckchem), 10 µM Forskolin (Sigma-Aldrich). For the first 2 days of culture 0.5 nM Wnt Surrogate (IPA Therapeutics) and 10 µM γ-27632 (Selleckchem) were added to CEM. Medium was refreshed every 2-3 days throughout the culture period. Intrahepatic cholangiocyte organoids (ICOs) were passaged every 5-7 days at a split ratio of 1:4-1:6. Cholangiocyte organoids were matured towards hepatocyte-like cells (ICO-Hep), as previously described (Huch., 2015). Briefly, cell were cultured in CEM + 25 ng/mL BMP7 (Peprotech) for three days. The media was then changed to hepatocyte differentiation medium (HDM) for 10 days. HDM consists of advanced DMEM/F12, supplemented with 1% (v/v) GlutaMax, 1% (v/v) Non-Essential Amino Acids, 1% (v/v) Penicillin-Streptomycin, 10 mM HEPES (all Gibco), 1% (v/v) B27 (minus vitamin A) (Thermo), 1% (v/v) N2 (Thermo), 10 nM gastrin (Sigma-Aldrich), 100 ng/mL FGF19 (Peprotech), 25 ng/mL BMP-7 (Peprotech) 50 ng/mL EGF (Peprotech), 25 ng/mL HGF (Peprotech), 0.5 µM A-83-01 (Selleckchem), 10 µM DAPT (Selleckchem) and 30 µM Dexamethasone (Sigma-Aldrich).

### RNA Isolation and qRT-PCR

Total RNA was isolated from organoids, tissue and primary cells using the RNeasy Mini Kit (QIAGEN) following the manufacturer’s instructions. In each case, the pellet was initially snap frozen and later resuspended in 350 µL of RLT lysis buffer solution containing 1% β-mercaptoethanol. The purified RNA was resuspended in 30 μL RNase-Free water. This was followed by reverse transcription and cDNA synthesis using iScript™ cDNA Synthesis Kit (Bio-Rad). The cDNA was then used for <u>qPCR</u> analysis with iTaq™ Universal SYBR®Green Supermix (Bio-rad). The samples were run on Bio-Rad CFX384 Touch Real-Time PCR systems (Bio-Rad) using Bio-Rad CFX Manager 1.1. To analyze gene expression of matured HBOs, organoids were matured in HMM for 4 days prior to harvesting. Gene expression was normalized to the mean of housekeeping genes (HMBS and RPL13a) and fold-change was set relative to donor matched PHH before culture.

### Total Protein Content Determination

Protein quantification using Pierce™ BCA Protein Assay Kit (Sigma-Aldrich) was carried out under manufacturer’s instructions. Briefly, 25 µL of each sample and standard together with 200 µL of the working reagent were pipetted into a microplate well and shaken for 30 seconds. The plate was covered and incubated up to 1 h at 37 °C, with Absorbance measured at 562 nm using a CLARIOstar® Plus microplate reader and MARS v4.00 software.

### EdU Incorporation

To evaluate EdU incorporation we used the Click-iT EdU Cell Proliferation Kit (Thermo). Briefly, spheroids or organoids were incubated with EdU overnight and then collected, washed and fixed in 4% paraformaldehyde for 45 minutes at 4°C. Section or wholemount staining was then performed according to the manufacturer’s instructions.

### Immunofluorescence Staining

Organoids were liberated from Matrigel with dispase and fixed in 4% paraformaldehyde for 45 minutes at 4°C. and then washed in 70% EtOH.

For sections, organoids were embedded into paraffin blocks, sections were cut, adhered to slides in 55°C oven, and hydrated. Depending on the antigen, slides were boiled in wither TE (pH 9.0) or citrate (pH 6.0) and rinsed in PBS. Organoids were permeabilized in PBS containing 0.1% Triton X-100 (PBT) for 15 minutes and then blocked using normal goat serum for 30 minutes. Primary antibodies were incubated for 1 hour at room temperature and then washed away with PBS. Secondary antibodies were incubated for 1 hour at room temperature and then washed away with PBS. Slides were then incubated with DAPI for 10 minutes, washed and then mounted for imaging.

For wholemount organoids were stained using adapted protocol of Dekkers *et al*.^46^. Briefly, organoids were permeabilized in PBT for 10 minutes, and then blocked in PBS containing 0.1% Triton X-100 and 0.2% BSA (OWB) for 10 minutes. Primary antibodies were incubated for 1 hour at room temperature and then washed away with OWB. Secondary antibodies were incubated for 1 hour at room temperature and then washed away with OWB. Organoids were then incubated with DAPI for 10 minutes and washed with PBS. Organoids were cleared using fructose-glycerol solution for 30 minutes and then imaged TCS SP8 X confocal microscope (Leica).

### Functional Analysis of Hepatobiliary Organoids and ICO-Heps

To evaluate glycogen storage, organoids were stained with periodic acid-Schiff staining kit (Sigma). LDL uptake was detected using Image-iT™ Low Density Lipoprotein Uptake Kit, pHrodo™ Red (Thermo). Albumin secretion was determined using human albumin ELISA kit (Bethyl Laboratories). For analysis of albumin secretion and LDL-uptake in HEM, hepatobiliary organoids were starved of serum for 48 hours before beginning the experiment. For analysis of albumin secretion in matured organoids, HBOs were matured in HMM for 4 days prior to beginning the experiment. For HBOs and ICO-Heps culture media was collected after 48 hours and human albumin was measured. For analysis of albumin secretion in 2D sandwich cultured PHHs (day 2-3) media was collected after 24 hours. All kits were used according to the manufacturer’s instructions.

The drug-metabolizing activity of HBOs and ICO-Heps was evaluated using a cocktail of parental compounds, as previously described^48^. These included 5 μM Midazolam (BUFA; metabolites: 1’-Hydroxymidazolam, Sigma-Aldrich; and 1’-Hydroxymidazolam glucuronide, LGC Standards), 15 μM Dextromethorphan (Santa Cruz Biotechnology; metabolite: Dextrorphan tartrate, Sigma-Aldrich), 20 μM Tolbutamide (Sigma-Aldrich; metabolite: 4-Hydroxytolbutamide, LGC Standards), 15 μM Phenacetine (Sigma-Aldrich; metabolite: Acetaminophen, Sigma-Aldrich), 20 μM Bupropion-HCl (Sigma-Aldrich; metabolite: Hydroxybupropion, Sigma-Aldrich) and 12 μM 7-Hydroxycoumarin (Umbelliferone, Sigma-Aldrich; metabolite: 7-Hydroxycoumarin glucuronide, LGC Standards). Cultures were exposed to the cocktail in HEM, HMM or ICO-DM in a humidified atmosphere at 37°C and 5% CO^2^ for 48 hours. 200 μl culture medium was collected in the 1.5-ml Short Thread Vials with the ND9 Short Thread Screw Caps (BGB Analytik), diluted 8 times with ultrapure MeOH, vortexed and stored at −20°C until the quantification of the parental compounds and metabolites using LC-MS/MS, as previously reported^47^. Briefly, prior to the analysis, samples were centrifuged for 10 min at 1500 g to precipitate any protein. Standards of the parental compounds and metabolites were prepared in the same medium (matrix) as samples. 1 μL solution was injected for measurements. Standards and samples were analyzed in a single run using a Shimadzu triple-quadrupole LCMS 8050 system with two Nexera XR LC-20AD pumps, a Nexera XR SIL-20AC autosampler, a CTO-20AC column oven and an FCV-20AH2 valve unit (Shimadzu). The compounds were separated on a Synergi Polar-RP column (150 × 2.0 mm, 4 µm, 80 Å) with a 4 × 2 mm C18 guard column (4 × 2 mm, Phenomenex). The mobile phase consisted of 0.1% (*v*/*v*) formic acid in ultrapure water (A) and 0.1% (*v*/*v*) formic acid in MeOH (B), and was set as 100% A (0-1 min), 100% to 5% A (1-8 min), 5% A (8-9 min), 5% to 100% A (9-9.5 min) and 100% A (9.5-12.5 min). The total run time was 12.5 min, and the flow rate was 0.2 mL/min. Peaks were integrated using the LabSolutions software (Shimadzu). For each compound, a limit of quantification (LOQ) was determined based on a standard curve and a limit of detection (LOD) was determined based on a noise signal. For each sample set, a dose (time 0 hours) and a blank (negative control) were measured. Amounts of metabolites were normalized to 1 µg protein. For analysis of drug metabolism in HEM, HBOs were starved of serum for 48 hours before beginning the experiment. For analysis of drug metabolism in matured organoids, HBOs were matured in HMM for 4 days prior to beginning the experiment.

### Bile Acid Transport

To visualize bile acid transport, organoids were incubated with CLF [1 µM] or CDFDA [1 µM] for 15 minutes at 37°C. CLF/CDFDA was then washed away and Matrigel was digested with dispase to release intact hepatobiliary organoids. Organoids were washed with PBS, plated in either HEM or HMM and immediately imaged on EVOS FL (Thermo). Bile acid transport inhibition experiments were performed as follows. Organoids were pre-treated with Troglitazone [50 µM] in HEM or HMM for 45-60 minutes at 37°C. Media was then changed to media containing CLF [1 µM] +/-Troglitazone [50 µM] and organoids were incubated for 15 minutes at 37°C. CLF was then washed away and Matrigel was digested with dispase to release intact hepatobiliary organoids. Organoids were washed with PBS, plated in either HEM or HMM and immediately imaged on EVOS FL (Thermo). Mean fluorescence intensity was determined using ImageJ software.

### Lipid Treatment and Neutral lipid staining

Hepatobiliary organoids in HEM were matured in HMM for 2 days. After 2 days of maturation, organoids were switched to HMM supplemented with 0.4 mM Palmitate-BSA (PA:BSA = 6:1), 0.2 mM Oleate-BSA (OA:BSA = 6:1), 5 mM Sodium acetate and 0.2 mM Glycerol. Organoids were incubated in the lipid cocktail for up to 12 days with media refreshments every 3-4 days. To stain neutral lipids, organoids were liberated from Matrigel using dispase, washed with PBS, and incubated with PBS containing 1 µg/mL BODIPY 493/503 for 15 minutes at 37°C. Organoids were then washed twice with PBS and immediately imaged on EVOS FL (Thermo).

### Engineered Virus-like Particles Production and Transduction of PHH

Production of engineered virus-like particles (eVLPs) was performed by adopting the previously described protocol^48^. First, Gesicle Producer 293T cells were plated at a density of 6×10^6^ cells per T75 flask. After 24 hours, cells were transfected with plasmids expressing VSV-G (4 μg; Addgene #12259), Gag-Pol (3.375 μg; Addgene #35614), Gag-ABE8e (1.125 μg; Addgene #181751), and gRNA (4.4 μg; Addgene #65777) using Lipofectamine 2000. The Supernatant was collected 48 hours after transfection, centrifuged at 500 × g for 1 min, and filtered through 0.45-μm PVDF filters. The particles were then concentrated by ultracentrifugation at 50,000 × g, 4°C, for 2 hours and resuspended in 100 μl cold PBS.

Transduction of eVLPs to PHH cells was done by resuspending 3 x 10^4^ cells in 900 μl HSM supplemented with 100 μl eVLP suspension. The cells were seeded in ULA microwells at a density of 50-75 cells per well, centrifuged at 100g for 5 minutes at room temperature, and incubated with eVLP for 72 hours at 37°C and subsequently harvested to isolate the genomic DNA (gDNA). Isolation of gDNA was performed using the Quick DNA microprep kit (Zymogen) according to the manufacturer’s protocol. The region of interest in the HBEGF locus was amplified using Q5 Hot Start High-Fidelity 2x Master Mix (NEB) with 32 cycles of 98°C for 10 sec, 59°C for 30 sec, and 72°C for 30 sec. PCR products were cleaned up using NucleoSpin Gel and PCR Clean-Up kit (Macherey-Nagel) and Sanger sequenced using the forward primer at Macrogen Europe. To assess gene editing efficiency, sequencing results were analyzed using EditR web application^49^.

### Preparation of Lipid nanoparticles (LNPs) and Transfection of PHH

LNPs were produced by microfluidics mixing in a NanoAssemblr Benchtop device (Precision Nanosystems, Vancouver, Canada). Before production, lipids were diluted to a total lipid concentration of 10 mM in pure ethanol while mRNA was diluted in a 100mM acetate buffer (pH 4). GFP mRNA was produced in-house using the Hiscribe® T7 Quick high yield RNA synthesis kit and 3’-O-me-m7G cap analog (New England Biolabs, MA, USA) following the manufacture’s protocol. LNPs were produced with a wt/wt ratio (ionizable lipid/mRNA) of 12:1 at a total flow rate of 9 mL/min, flow rate ratio of 3:1 (aqueous: lipid phase) at room temperature. LNPs were composed of ionizable lipid/Cholesterol (Sigma-Aldrich, Saint Louis, MO, USA)/DSPC (Lipoid, Ludwigshafen am Rhein, Germany)/DMG-PEG2000 (Avanti polar lipids, Inc, Alabaster, AL, USA). at a molar ratio of 50/38.5/10/1.5 respectively. Four different formulations were made using the following ionizable lipids: DLin-MC3-DMA (The University of British Columbia, Vancouver, Canada), SM-102, ALC-0315 and Lipid 5 (Cayman Chemicals, Ann Arbor, MI, USA). Also 2 additional formulations were made using SM-102 and DLin-MC3-DMA, but cholesterol was substituted by b-sitosterol. LNPs were then dialyzed overnight into TBS using slide-a-LyzerTM dialysis cassettes G2 20kDa (Thermo). LNPs were characterized by size, zeta potential and encapsulation efficiency.

Transfection of LNPs to PHH cells was done by resuspending 2 x 10^4^ cells in HSM supplemented with either 50, 100, 250 or 500 ng of LNPs. The cells were seeded in ULA microwells at a density of 50-75 cells per well, centrifuged at 100g for 5 minutes at room temperature and incubated at 37°C for 72 hours and subsequently imaged on EVOS FL (Thermo). Mean fluorescence intensity was determined using ImageJ software.

## Supplemental Information

### Supplemental Figures

**Figure S1.**
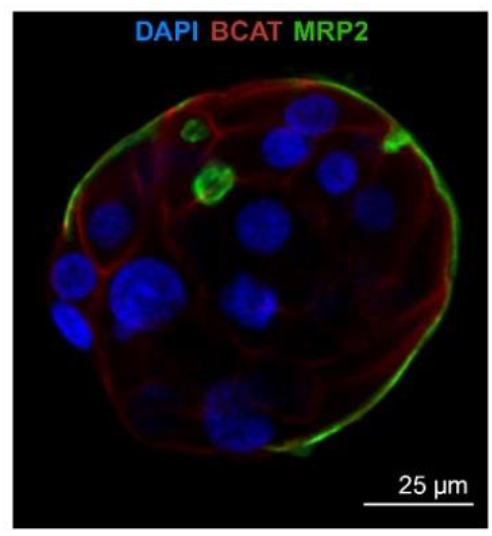
MRP2 localization in Hepatocyte Spheroids. Confocal single-plane image of BCAT (red) and MRP2 (green) in hepatocyte spheroids in HSM, scale bar = 25 µm.

**Figure S2.**
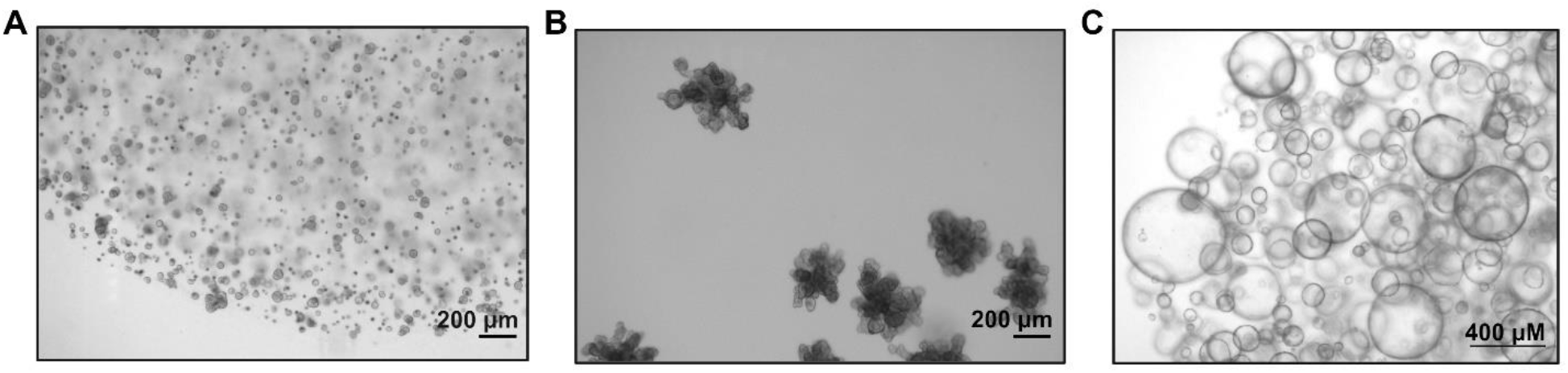
Hepatocyte Organoids, Hepatobiliary Organoids and Intrahepatic Cholangiocyte Organoids. A) Brightfield image of hepatocyte organoids (day 15 post seeding), scale bar = 200 µm. B) Brightfield image of hepatobiliary organoid (day 15 post seeding), scale bar = 200 µm. C) Brightfield image of intrahepatic cholangiocyte organoids (passage 4), scale bar = 400 µm. All images from single donor (age 27).

**Figure S3.**
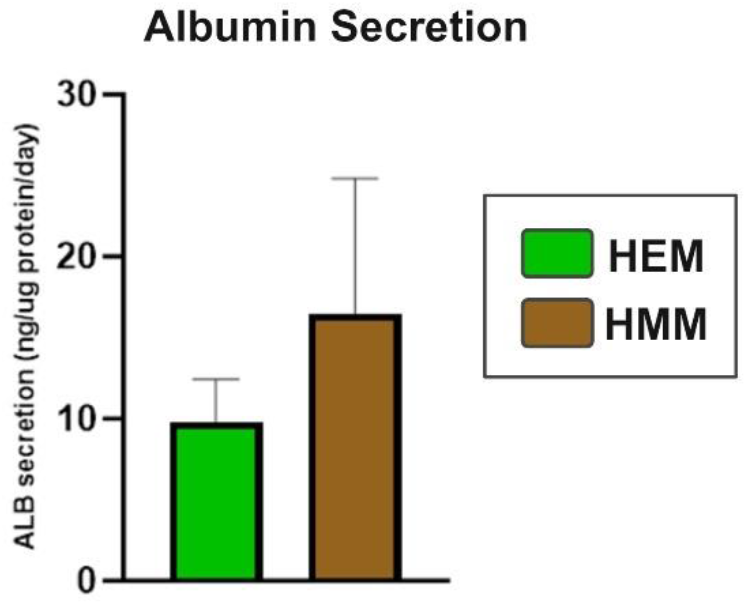
Albumin secretion of Hepatobiliary Organoids. Albumin secretion of HBOs after 48 hours in HEM and HMM. Graph presents mean results of 3 replicates from 3 individual wells of a single donor. Results are presented as ng albumin/ µg protein/day.

**Figure S4.**
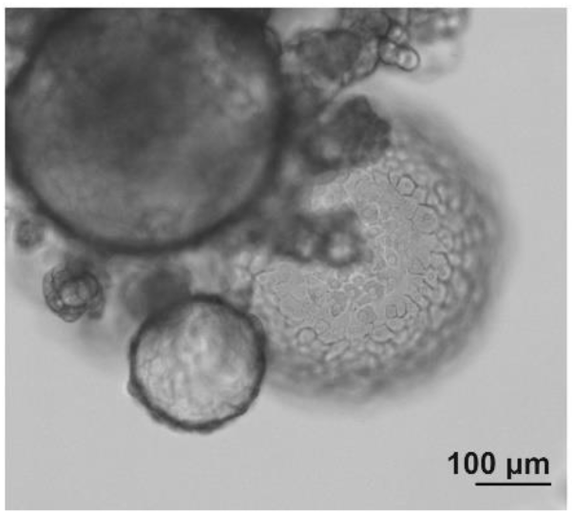
HBOs after single-cell passaging. Passage 4, scale bar = 100 µm

## Supplementary Tables

**Table S1.**
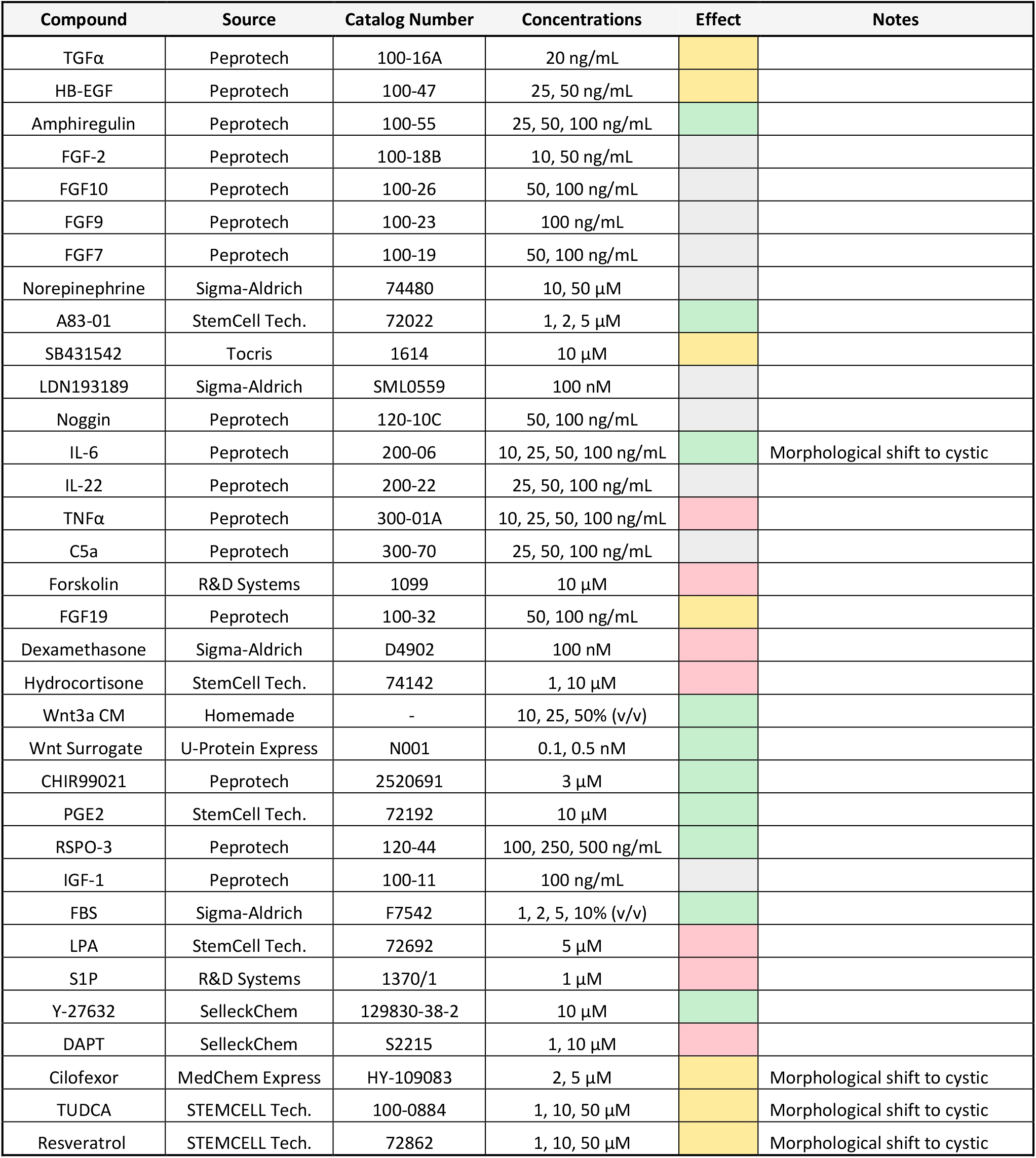
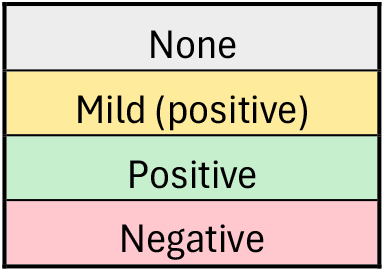
Compounds Tested for ability to promote hepatocyte proliferation on top of EGF, HGF, Nicotinamide, N-acetylcysteine and Gastrin

| Compound | Source | Catalog Number | Concentrations | Effect | Notes |
| --- | --- | --- | --- | --- | --- |
| TGFα | Peprotech | 100-16A | 20 ng/mL | | |
| HB-EGF | Peprotech | 100-47 | 25, 50 ng/mL |  |  |
| Amphiregulin | Peprotech | 100-55 | 25, 50, 100 ng/mL |  |  |
| FGF-2 | Peprotech | 100-18B | 10, 50 ng/mL |  |  |
| FGF10 | Peprotech | 100-26 | 50, 100 ng/mL |  |  |
| FGF9 | Peprotech | 100-23 | 100 ng/mL |  |  |
| FGF7 | Peprotech | 100-19 | 50, 100 ng/mL |  |  |
| Norepinephrine | Sigma-Aldrich | 74480 | 10, 50 μM | | |
| A83-01 | StemCell Tech. | 72022 | 1, 2, 5 μM | | |
| SB431542 | Tocris | 1614 | 10 μM | | |
| LDN193189 | Sigma-Aldrich | SML0559 | 100 nM |  |  |
| Noggin | Peprotech | 120-10C | 50, 100 ng/mL |  |  |
| IL-6 | Peprotech | 200-06 | 10, 25, 50, 100 ng/mL |  | Morphological shift to cystic |
| IL-22 | Peprotech | 200-22 | 25, 50, 100 ng/mL |  |  |
| TNFα | Peprotech | 300-01A | 10, 25, 50, 100 ng/mL | | |
| C5a | Peprotech | 300-70 | 25, 50, 100 ng/mL |  |  |
| Forskolin | R&D Systems | 1099 | 10 μM | | |
| FGF19 | Peprotech | 100-32 | 50, 100 ng/mL |  |  |
| Dexamethasone | Sigma-Aldrich | D4902 | 100 nM |  |  |
| Hydrocortisone | StemCell Tech. | 74142 | 1, 10 μM | | |
| Wnt3a CM | Homemade | - | 10, 25, 50% (v/v) |  |  |
| Wnt Surrogate | U-Protein Express | N001 | 0.1, 0.5 nM |  |  |
| CHIR99021 | Peprotech | 2520691 | 3 μM | | |
| PGE2 | StemCell Tech. | 72192 | 10 μM | | |
| RSPO-3 | Peprotech | 120-44 | 100, 250, 500 ng/mL |  |  |
| IGF-1 | Peprotech | 100-11 | 100 ng/mL |  |  |
| FBS | Sigma-Aldrich | F7542 | 1, 2, 5, 10% (v/v) |  |  |
| LPA | StemCell Tech. | 72692 | 5 μM | | |
| S1P | R&D Systems | 1370/1 | 1 μM | | |
| Y-27632 | SelleckChem | 129830-38-2 | 10 μM | | |
| DAPT | SelleckChem | S2215 | 1, 10 μM | | |
| Cilofexor | MedChem Express | HY-109083 | 2, 5 μM | | Morphological shift to cystic |
| TUDCA | STEMCELL Tech. | 100-0884 | 1, 10, 50 μM | | Morphological shift to cystic |
| Resveratrol | STEMCELL Tech. | 72862 | 1, 10, 50 μM | | Morphological shift to cystic |

|  |
| --- |
| None |
| Mild (positive) |
| Positive |
| Negative |

**Table S2.** Age and Sex and Culture Time of Primary Human Hepatocytes Donors.

| Donor | Age | Sex | Cultured as HBOs |
| --- | --- | --- | --- |
| 1 | 0 | Male | 3 months |
| 2 | 16 | Male | 1 month+* |
| 3 | 24 | Male | 1 month+* |
| 4 | 27 | Female | 3 months |
| 5 | 47 | Male | - |
| 6 | 50 | Female | 2 months |
| 7 | 51 | Male | - |
| 8 | 60 | Female | 2.5 months |
| 9 | 70 | Female | 1.5 months |

\*To be determined
- Not successful

## Notes

### Competing Interest Statement

The authors have declared no competing interest.

